# The microRNA156/*SPL9* module mediates auxin response to facilitate apical hook maintenance in *Arabidopsis*

**DOI:** 10.1101/2024.07.16.603710

**Authors:** Flaviani G. Pierdona, Ana Julia de Moraes Silva, Mateus Henrique Vicente, Laura Taylor, Ullas Pedmale, Fabio T. S. Nogueira

## Abstract

Auxin coordinates cell growth by promoting or inhibiting cell expansion during etiolated seedling development, but whether and how microRNA modules participate in this process remains unclear. Here, we show the miRNA156/*SQUAMOSA PROMOTER-BINDING-PROTEIN-LIKE9* (miR156/*SPL9*) module is critical for skotomorphogenesis. Perturbation of the miR156/*SPL9* module affected skotomorphogenesis, as the loss of miR156 function or *SPL9* de-repression led to shorter hypocotyl, higher hook angle, and delayed hook opening. Opposing phenotypes were observed in dark-grown *spl9* and miR156-overexpressing seedlings. Importantly, loss of miR156-dependent *SPL9* regulation triggered apical hook formation even under reduced levels of endogenous auxin. miR156-targeted *SPL9* arrested cell expansion by repressing *small auxin-up RNA19* (*SAUR19*) gene in a *FRUITFULL* (*FUL*)-dependent and independent manner. The conserved miR156/*SPL9/15* module also affects skotomorphogenesis in tomato, impacting its successful soil emergence. Our findings unravel how the miR156/*SPL9* module plays a pivotal role in the auxin network coordinating apical hook development to enable appropriate seedling emergence.

## INTRODUCTION

Seedling developmental plasticity in response to the environment is essential for plant establishment^1^. Seeds are sown under the soil, where it is protected from predators and extreme environment conditions. The absence of light, in this condition, triggers skotomorphogenesis, a developmental process that protects and promotes faster seedling emergence. Understanding the molecular mechanisms that control the early stages of development and seedling emergence has become essential to obtain better seedling vigor and stand establishment in crop production^2^. During soil emergence, the energy resource is allocated to fast hypocotyl growth, instead of roots or cotyledons, its upper part forms a simple curved structure, namely the apical hook, which is transient but crucial to protect the shoot apical meristem (SAM) and cotyledons^3^. Apical hook development may represent the most intensively investigated aspect of skotomorphogenesis since Darwin’s days^4^. Importantly, hook development is a dynamic model system which can be used to address questions relating to differential cell growth and seedling morphogenesis response in transition from dark to light condition^5,6^.

Apical hook development is divided into three main phases: formation, maintenance, and opening, each of these phases is marked by a differential cell expansion rate on both sides of the hypocotyl, regulated by a complex network, including light signaling and hormones^5^. Among the hormones ethylene and auxin are central for skotomorphogenesis regulation, especially in apical hook development^7^. Exogenous ethylene treatment triggers a triple response in dark-grown seedlings, which exhibit short roots and hypocotyls, as well as an exacerbated apical hook^8^. Ethylene-insensitive mutant analyses revealed the transcription factor *ETHYLENE-INSENSITIVE PROTEIN2* (*EIN2*) as the central component of ethylene signaling, including in apical hook formation^9^. Moreover, ethylene is an essential activator of auxin transport, and, in turn, the *AUXIN RESISTANT 1* (*AUX1*) is necessary for ethylene-based apical hook regulation^10^. Meanwhile, during the hook formation phase, intercellular auxin transport machinery establishes an auxin signaling maximum at the inner side of the hook, in which higher concentrations of auxin asymmetrically repress growth, leading to the curvature of the hypocotyl and subsequent apical hook formation^11^. Auxin promotes cell elongation in the hypocotyl through an acid-growth mechanism, but high levels of this hormone repress the expression of some *auxin-induced small auxin-up RNA* (*SAUR*), which leads an increase in PP2C.D (D clade type 2C protein phosphatase) activity and consequently inhibition of the plasma membrane proton pump (PM H+-ATPase), reducing cell elongation^12^. Members of the *SAUR* family are also target by the MADS box transcription factor AGAMOUS-LIKE8 (AGL8)/FRUITFULL (FUL), inhibiting early apical hook opening^13^. However, how auxin-dependent regulation of distinct *SAUR* genes is orchestrated in the apical hook an whether it is mediated by FUL is still unclear^12^.

Circumstantial evidence suggests that microRNAs (miRNAs) may also contribute to the control of kotomorphogenesis^14,15^. miRNAs are endogenous noncoding RNAs (20 to 24 nt) that regulate several developmental processes and environmental responses^16^. Long primary transcripts (pri-miRNA) are transcribed from *MIRNA* genes and then processed into miRNA duplexes by the RNAase III family enzyme DICER-LIKE1 (DCL1) along with two accessory proteins, the dsRNA-binding protein HYPONASTIC LEAVES 1 (HYL1) and the C2H2 zinc finger protein SERRATE (SE). The resulting miRNA duplex is loaded into ARGONAUTE1 (AGO1), in which one strand of the duplex functions as the miRNA that guides AGO1 to the target messenger RNA based on sequence complementarity, resulting in the degradation or translational repression of the target transcripts^16^.

Mutations in miRNA biogenesis core enzymes, such as SE, DCL1, and AGO1 lead to shorter hypocotyls and more closed apical hooks, probably through the combined action of specific miRNAs and auxin^14,17^. Indeed, the auxin gradient is altered in the *se-1* apical hook, in line with its prolonged hook maintenance phase^18^, suggesting miRNA-dependent auxin accumulation during skotomorphogenesis. Small RNA sequencing in soybean reveals that miRNA transcript levels increased in the apical hook after treating dark-grown seedlings with far-red light. Hook opening of *Arabidopsis ago1* seedlings is reduced and delayed in response to far-red light^19^. Several miRNA-mRNA pairs were found to be important regulators of dark-to-light transitions. For instance, degradome signatures in de-etiolated *Arabidopsis* seedlings revealed that light treatment induces higher degradation of the targets of the miR156/miR157 family^20^. The miR157 was described as a negative regulator of photomorphogenesis by directly targeting the transcription factor *ELONGATED HYPOCOTYL5* (*HY5*)^15^. The miR156 is differentially expressed in dark and light-grown seedlings^14^, but its role in skotomorphogenesis is unknown.

miR156 targets members of the *SQUAMOSA PROMOTER BINDING PROTEIN-LIKE (SPL)* family^21,22^ establishing an age-dependent pathway to control phase transition^23^ as well as different aspects of plant development^24–26^. The miR156/*SPL* module has been also shown to be an important regulator of environmental responses^27–29^. In adult *Arabidopsis* plants, for instance, *MIR156* genes are repressed by PIFs to induce shade avoidance through the *SPL*s^27^. In seedlings, miR156 releases high temperature-mediated morphogenesis by reducing auxin sensitivity^28^. Indeed, the miR156/*SPL* module regulates auxin responses in different aspects of development. miR156-based repression of *SPL10* coordinates auxin transport in the *Arabidopsis* root meristem affecting primary root growth^25^. In tomato, the overexpression of miR156-targeted *SlSBP15* represses axillary bud development by inhibiting auxin transport^30^. In addition, *FUL*, a direct downstream target of miR156-targeted *SPLs*^31^, has been shown to be a key repressor of auxin-responsive genes during apical hook development^13^. However, whether the miR156-*SPL*-*FUL* genetic circuity operates in the auxin-based apical hook development remains unclear.

To unravel the role of miRNAs, and more specifically the miR156/*SPL* module during skotomorphogenesis, here we used apical hook as a model to test whether perturbations in this miRNA module affect hook development. In dark-grown seedlings, the miR156-based regulation of *SPL9* is fundamental for proper apical hook formation and maintenance. miR156 locally represses *SPL9* expression in the inner part of the apical hook, which in turn represses cell growth by modulating auxin responses and promoting hypocotyl curling. Importantly, we propose an auxin-based mechanism of cell expansion in which SPL9 regulates *FUL* and *SAUR19* in etiolated seedlings. We showed that the miR156/*SPL* role in the skotomorphogenesis is conserved in tomato and impacts seedling emergence rate and stand establishment. We provide evidence of a new role for the miR156/*SPL9* module in this crucial early developmental process that facilitates seedling emergence.

## RESULTS

### The miR156/*SPL* module regulates *Arabidopsis* apical hook development during skotomorphogenesis

Dark-grown *se-1* mutant exhibits low levels of the miR156 and shorter hypocotyls as well as more closed apical hooks^17^. Because some of the miR156-dependent developmental processes in juvenile plants overlap with the SE-mediated responses, and the miR156 overexpression partially rescues *se-1* mutant phenotypes^32^, we hypothesized that miR156 and its targets are one of the main players in the miRNA-dependent regulation of skotomorphogenesis. To test this conjecture, we evaluated the apical hook and hypocotyl development of dark-grown seedlings (Fig. 1a, b) overexpressing the miR156 (miR156oe)^23,33^. Three-day-old miR156oe seedlings exhibited a significant decrease in the apical hook curve and longer hypocotyls compared to Col-0 grown under dark conditions (Fig. 1c, d). On the other hand, the loss of function of four *MIR156* genes (*mir156a/b/c/d*) resulted in the opposite phenotype, i.e., more intense apical hook and smaller hypocotyl (Fig. 1c, d). These phenotypes are consistent with those of *se-1* mutants^18^, and support the hypothesis that miR156 is one of the crucial miRNAs regulating cell expansion during skotomorphogenesis. We then examined hook angle in response to light in miR156oe and *mir156a/b/c/d* seedlings, and in a transgenic line constitutively expressing a miR156 target mimicry (MIM156), in which miR156 activity is blocked^34^. While hook opening was repressed in both *mir156a/b/c/d* and MIM156 seedlings, resembling *se-1* seedlings^18^, miR156oe hook opening showed an opposite phenotype compared to Col-0 (Supplementary Fig. 1a). Because the miR156/*SPL* module regulates hypocotyl elongation at warm temperatures^28^, we also evaluated hypocotyl elongation in the dark to light transition. Both dark-grown *mir156a/b/c/d* and MIM156 seedlings displayed shorter hypocotyls, whereas miR156oe exhibited larger hypocotyls compared to Col-0 seedlings (Supplementary Fig. 1b). Together, these results strongly suggest that the miR156 is essential to control the transition from skotomorphogenesis to photomorphogenesis.

To better understand the role of miR156 in skotomorphogenesis, we focused on the apical hook development, a model to study differential cell elongation and transition from skotomorphogenesis to photomorphogenesis^5,35^. We then analyzed the miR156 spatial expression in the apical hook by *GUS* reporter gene. Because *MIR156A* and *MIR156C* genes are the most functionally significant *MIR156* genes in vegetative development ^36–38^, we selected *MIR156A-GUS* and *MIR156C-GUS* lines ^38^. Both *MIR156A-GUS* and *MIR156C-GUS* reporter genes were detected at the apical region of dark-grown seedlings, especially in the hook (Fig. 1e). In addition, the spatial miR156 activity was examined in response to light employing sensor lines constitutively expressing *GFP-N7* containing the *SPL3* 3’UTR harboring the miR156 recognition site (GFP-miR156 sensor), or *GFP-N7* containing the *NOS* terminator (control sensor)^39^. Sensor analyses revealed high activity of miR156 in the hook, indicated by greater levels of repression of *GFP* expression at the apical region of the seedlings growing under dark. This observation was intensified under light conditions (Supplementary Fig. 1c).

Given that miR156 targets 10 *Arabidopsis SPL*s^40^, we asked whether miR156 mediates hook formation by repressing the *SPLs*. Considering that all ten miR156-targeted *SPL*s are classified into three functionally distinct groups^41^, we selected *SPLs* representing each group: *SPL9, SPL10, SPL15* (group 1), *SPL3* (group 2), *SPL6* (group 3). Group 1 is the largest ^41^, thus we selected more than one *SPL* gene. To examine the functions of *SPLs* in the miR156-mediated control of hook formation, we analyzed apical hook in 3-day-old dark-grown seedlings expressing miR156-resistant (*rSPL*) version of the *SPLs*^41^. Among the rSPLs evaluated, the de-repression of *SPL9* in rSPL9 seedlings resulted in stronger curvature of the apical hook compared to Col-0, similar to rSPL15 (Fig. 1f, g). Despite dark-grown rSPL3, rSPL6 and rSPL10 seedlings showed an intermediate phenotype compared to Col-0 and other rSPLs, we did not observe a significant difference in hook angle of these genotypes (Fig. 1g).

Because *SPL9* and *SPL15* show redundant functions in the juvenile-to-adult vegetative transition^32^, and our results indicated a similar phenotype, we focused mostly on *SPL9* role in hook formation. To compare the expression patterns among the *rSPLs*, we employed rSPL version of the *SPLs* fused to the reporter *GUS* ^41^. *rSPL3-GUS* was lowly expressed at the apex, whereas *rSPL6-GUS* and *rSPL10-GUS* were faintly or not expressed (Fig. 1h). On the other hand, rSPL9-GUS seedlings showed strong *GUS* signal at the apex (Fig. 1h). In addition, we compared hook phenotype among Col-0, rSPL9-GUS, and a miR156-sensitive *SPL9* fused to *GUS* (sSPL9-GUS) line. As expected, 3-day-old dark-grown rSPL9-GUS seedlings displayed higher hook angle, whereas sSPL9-GUS and Col-0 seedlings exhibited similar hook phenotypes (Supplementary Fig. 2a). Moreover, we did not detect *GUS* signal at the apex of dark-grown sSPL9-GUS seedlings, indicating that *SPL9* is strongly repressed by miR156 at this developmental stage (Supplementary Fig. 2b). *sSPL9-GUS* was detected in young leaves of 10-day-old seedlings grown under light condition (Supplementary Fig. 2c). Dark-grown *spl9-4* and *spl9/15* mutants displayed more opened hooks compared to Col-0, but much less than miR156oe seedlings (Supplementary Fig. 3a). This observation suggests that, in addition to *SPL9*, other miR156-targeted *SPLs* may be associated with hook formation. Interestingly, the quintuple mutant of *SPLs* belonging to group I (*spl2/9/11/13/15* or *q-spl* ^41^) displayed hook phenotype similar to *spl9-4* and *spl9/15* when growing in the dark (Supplementary Fig. 3a), reinforcing the crucial role of *SPL9* (and likely *SPL15*) in the regulatory network orchestrating apical hook formation. Because auxin asymmetrically accumulates at the hook to inhibit cell elongation and drive hook formation^7^, we examined rSPL9 localization in 3-day-old dark-grown seedlings expressing the *SPL9pro::GFP-rSPL9* construct (GFP-rSPL9). GFP-rSPL9 specifically accumulated in the nuclei of cells on the concave side of the hook (Fig. 1i), which coincides with auxin maxima^7^. Collectively, our observations indicated that miR156 regulates apical hook development mainly by repressing *SPL9* activity.

**Fig. 1.**
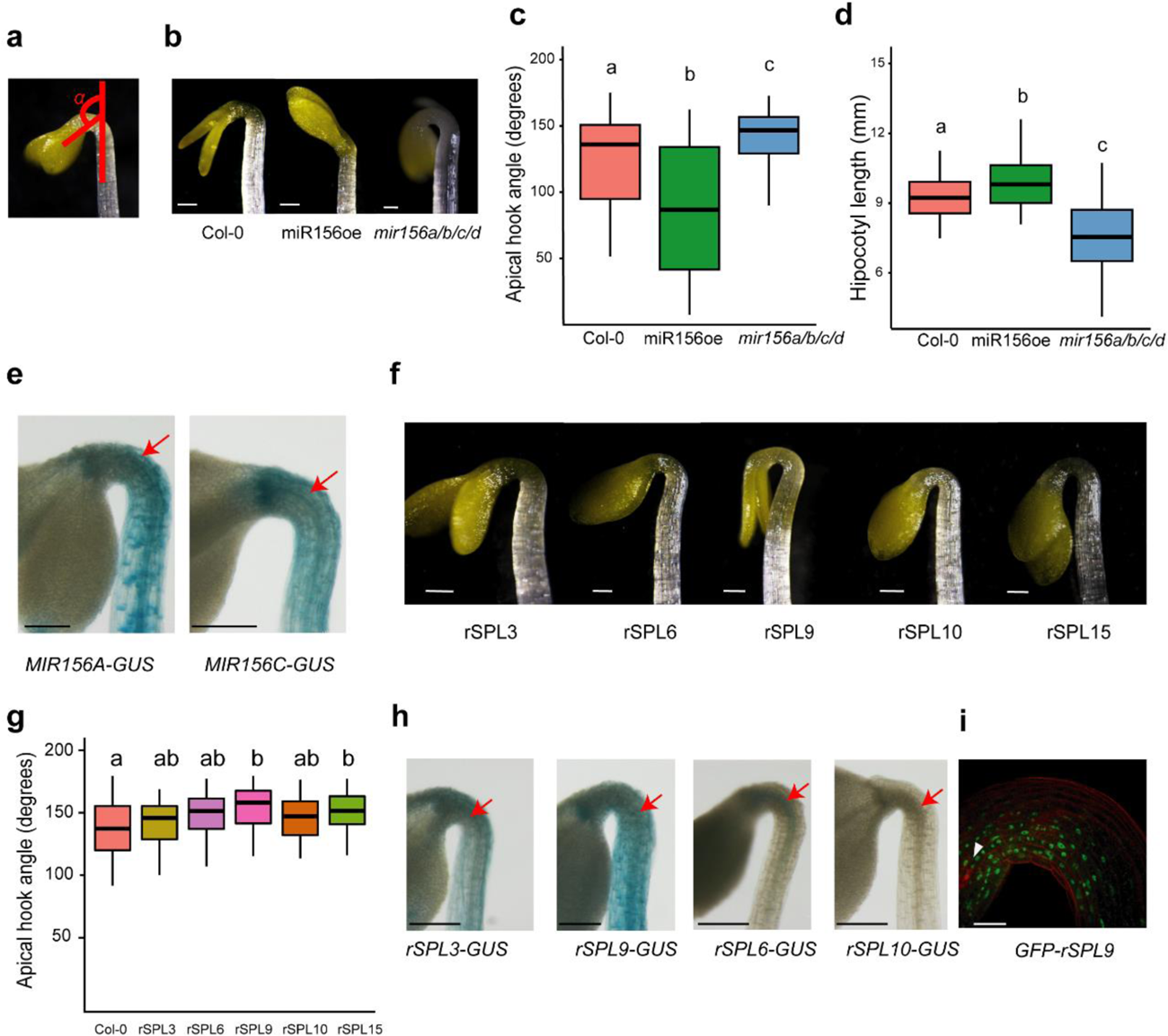
The miR156/*SPL* module regulates *Arabidopsis* apical hook development. **a** Scheme of the complementary angle (α) between the cotyledons and hypocotyl. **b** Representative images of 3-day-old *Arabidopsis* seedlings grown in darkness. Wild type (Col-0), miRNA156 overexpression (miR156oe), and quadruple loss of miR156 function mutant (*mir156a/b/c/d*). **c** Angle of the apical hook of 3-day-old dark-grown Col-0, miR156oe, and *mir156a/b/c/d*. Different letters mean statistical difference by ANOVA followed by Tukey test, 5% significance. **d** Hypocotyl length of 3-day-old dark-grown Col-0, miR156oe, and *mir156a/b/c/d*. Different letters mean statistical difference by ANOVA followed by Tukey test, 5% significance. **e** Expression pattern of *MIR156A* and *MIR156C* precursors fused to *GUS* (*proMIR156A::MIR156A-GUS* and *proMIR156C::MIR156C-GUS*) in 3-day-old dark-grown seedlings. Scale bar: 100 µm. **f** Representative images of 3-day-old *Arabidopsis* seedlings grown in darkness. miR156-resistant version (r) of *SPL3*, *SPL6*, *SPL9*, *SPL10*, and *SPL15*. **g** Apical hook angle of 3-day-old dark-grown Col-0 and miR156-resistant *SPL* versions-expressing (rSPL3, rSPL6 rSPL9, rSPL10, and rSPL15) seedlings. Different letters mean statistical difference by ANOVA followed by Tukey test, 1% significance. **h** Representative images of dark-grown, 3-day-old seedlings expressing *rSPLs* fused to *GUS* (*rSPL3-GUS, rSPL9-GUS, rSPL6-GUS,* and *rSPL10-GUS*) driven by their own promoters. Scale bar: 100 µm. **i** Representative image of dark-grown, 3-day-old dark-grown seedling expressing GFP-rSPL9 (*SPL9pro::GFP-rSPL9*) in the apical hook. White triangle indicates the cotyledons direction. Scale bar: 50 µm.

### The miR156-targeted *SPL9* is sufficient to control hook formation and maintenance phases

During the maintenance phase, the apical hook remains closed until light signal or time induces its opening^5^. To further understand the miR156-targeted *SPL9* function in the hook regulatory network, we generated transgenic lines expressing *GUS* driven by the *SPL9* promoter (*SPL9pro::GUS*). We then compared *SPL9* promoter activity with rSPL9-GUS accumulation (*SPL9pro::rSPL9-GUS* line) during formation, maintenance, and opening phases (Fig. 2a, b). *GUS* signal was stronger during the formation and maintenance phases, in both *SPL9pro::GUS* and *SPL9pro::rSPL9-GUS* lines (Fig. 2a, b). In addition, *GUS* signal from *SPL9pro::GUS* was detected only in the cells of the concave side (Fig. 2a, arrow), similar to the localization of GFP-rSPL9 protein (Fig. 1i). These observations reinforce that *SPL9* is expressed in the inner side of the arch, and it is regulated by miR156 mostly in the initial phases of hook development.

We next examined hook angle in response to light in miR156oe, rSPL9, and *spl9/15* seedlings compared to Col-0. As expected, lower expression of *SPL*s (including *SPL9*) in the miR156oe led to faster hook opening, similar to *spl9/15*. On the other hand, de-repression of *SPL9* in rSPL9 seedlings repressed apical hook opening in response to light (Supplementary Fig. 4). This observation supports the conclusion that miR156-based *SPL9* regulation is required for seedlings at dark to light transition. To corroborate this conclusion, we employed *Arabidopsis* lines expressing *rSPL9* fused to a glucocorticoid receptor (rSPL9-GR), in which we were able to modulate rSPL9 activity by adding the glucocorticoid Dexamethasone (DEX). rSPL9-GR was activated in the formation/maintenance phases by adding DEX in the germination medium, and at 3 days, followed by light treatment to induce the opening phase. As expected, in the absence of DEX, the rSPL9-GR shows a wild type phenotyoe (Fig. 2c), oppositly, in prescence of DEX, the rSPL9-GR present the same phenotype as rSPL9 (Fig. 2d). Interestingly, *SPL9* de-repression increased apical hook angle, but did not affected the apical hook oppening when rSPL9-GR was activated only at the formation/maintenance phases (Fig. 2e). DEX treatment at oppening phase, promoting a late activation of rSPL9 is also not sufficient to inhibit apical hook oppening (Fig. 2f). Together, our results strongly support the conclusion that miR156 is required for regulating *SPL9* in the concave cells to promote apical hook formation and maintenance phases.

We then asked whether miR156-targeted *SPL9* controls the differential cell expansion by measuring the cell size ratio between the concave and convex sides of the hook region from dark-grown Col-0, rSPL9, and *spl9-4* seedlings during maintenance phase (Fig. 2d, e). The average cell size from the convex side of rSPL9 apical hook is 2.4-fold larger than the concave cells, which was significantly higher than Col-0 (1.8-fold) and *spl9-*4 (1.6-fold) ratios (Fig. 2e). These observations were in line with the more intense apical hook phenotype observed in dark-grown rSPL9 seedlings (Fig. 1). Both concave and convex epidermal cells of the rSPL9 hook were shorter than Col-0 (Fig. 2e), which might indicate additional roles of the miR156/*SPL9* module in coordinating hypocotyl cell expansion during etiolated seedling development. The loss of *SPL9* function in *spl9-4* seedlings did not significantly affect cell expansion (Fig. 2 d,e), reinforcing the importance of the miR156-based *SPL9* regulation of cell size during apical hook establishment and maintenance.

**Fig. 2.**
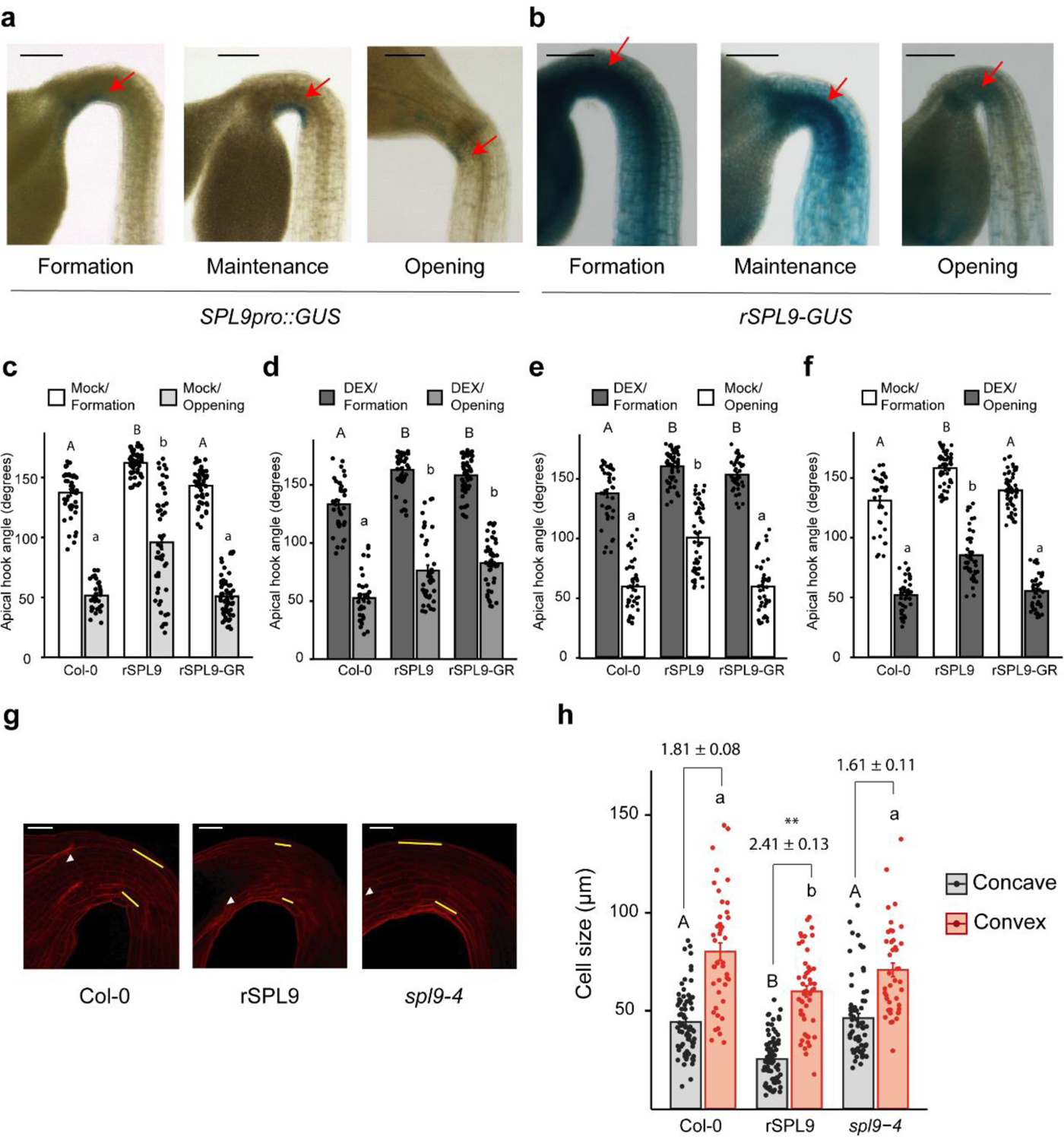
Perturbation of the miR156/*SPL9* module affects apical hook formation and maintenance. **a** *SPL9pro::GUS* expression in the apical hook formation (2-day-old), maintenance (3-day-old), and opening (4-day-old). Scale bar: 100 µm. **b** *SPL9pro::rSPL9-GUS* expression in the apical hook formation, maintenance, and opening. Scale bar: 100 µm. **c-f** Angle of the apical hook of wild type (Col-0), miRNA156-resistant *SPL9* version (rSPL9), and miRNA156-resistant *SPL9* version fused to the glucocorticoid receptor (rSPL9-GR). 0.1% DMSO (Mock) or 10 µM Dexamethasone (DEX) were added to de medium before seed germination (formation) and after 3 days in dark were treated with mock or DEX for three hours under white light (opening). Black dots represent individual data points. Different uppercase letters mean statistical significance among genotypes treated in formation phase; different lowercase letters mean statistical significance among genotypes after treatment in opening phase. ANOVA followed by Tukey test, 1% significance. **g** Representative confocal images of 3-day-old dark-grown seedlings of Col-0, rSPL9, and loss of *SPL9* function (*spl9-4*) grown in darkness and subsequently treated with propidium iodide. White triangle indicates the cotyledons direction. Yellow lines indicate a representative epidermal cell. Scale bar: 50 µm. **h** Measurement of the cell size in concave and convex sides of the apical hook from seedlings in **(g)**. Different uppercase letters mean statistical significance among genotypes for the concave cell size. Different lowercase letters mean statistical significance among genotypes for the convex cell size. ANOVA followed by Tukey test, 1% significance. Numbers above the bars represent the relative size of convex to concave epidermal cell ratio (mean ± SE., *n* = 8 biological replicates). **: statistically significant difference of the relative cell size compared to Col-0, two-tailed t-Test, 1% significance.

### The miR156/*SPL9* module intermediates auxin-based regulation of cell expansion in the apical hook maintenance phase

Our results demonstrated that *SPL9* controls the expansion of the inner cells of the apical hook (Fig. 2e). As the miR156/*SPL9* module regulates hypocotyl elongation at warm temperatures by modulating auxin sensitivity^28^, we hypothesized that this miRNA module is part of the auxin regulatory network controlling hook development. In the absence of light, ethylene induces the formation of auxin maxima, which in turn, plays an essential role in repressing cell elongation in the inner cells of the apical hook during formation and maintenance phases^12^. It was also recently reported the intrinsic connection between the miR156/*SPL* module and ethylene pathway in blueberry fruits^42^. Given that ethylene controls auxin differential distribution^10^, we asked whether ethylene may attenuate *MIR156* expression in the hook. Indeed, *MIR156C-GUS* expression was reduced at the apical region of dark-grown seedlings treated with the ethylene precursor 1-Aminocyclopropane-1-carboxylate (ACC) (Fig. 3a). Similarly, the synthetic auxin picloram repressed *MIR156C-GUS* expression at the apical region of dark-grown seedlings (Fig. 3a). Collectively, our results suggest that miR156 is part of the auxin response network that modulates apical hook formation.

**Fig. 3.**
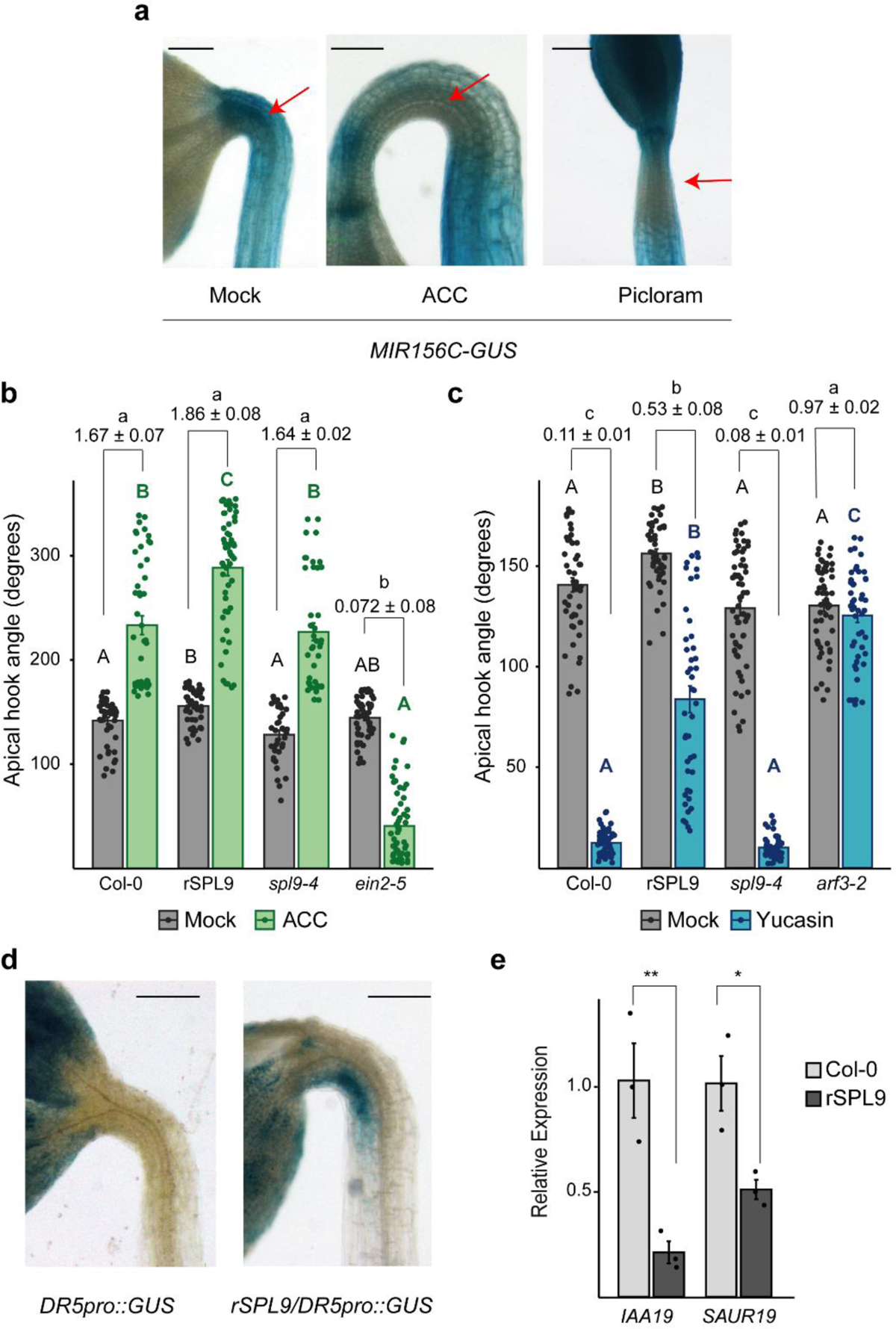
miR156-based repression of *SPL9* is necessary for auxin responses in the apical hook. **a** Expression of *MIR156C-GUS* in seedlings grown in 0.1% DMSO (Mock); 10 µM of ACC; or 5 µM of Picloram under dark conditions. scale bar: 100 µm. **b-c** Apical hook angle of 3-day-old dark-grown seedlings of Col-0, miR156-resistant *SPL9* version (rSPL9), the loss of *SPL9* function mutant (*spl9-4*), and the loss of *EIN2* function mutant (*ein2-5*) in **b** or the loss of *ARF3* function mutant (*arf3-2*) in **c**. Seeds were germinated in 0.1% DMSO (Mock) (gray bars); 10 µM of ACC (green bars in **b**) or 50 μM of the auxin inhibitor Yucasin (blue bars in **c**) Colored dots represent individual data points. Numbers above the bars represent the relative response of the hook angle, mean ± SE of the fold-changes between treatments and their respective mock, *n* = 3 biological replicates in each genotype. Different black uppercase letters mean statistical significance between the genotypes in mock. Different colored uppercase letters mean statistical significance between the genotypes in ACC (green) or Yucasin (blue). Different lowercase letters mean statistical significance between the relative response of each genotype. ANOVA folloed by Tukey test, 5% significance. **d** *DR5pro::GUS* expression pattern in *DR5pro::GUS* and double *DR5pro::GUS*/rSPL9 lines. Scale bar: 100 µm. **e** Relative expression of *IAA19* and *SAUR19* in apices of dark-grown Col-0 and rSPL9 seedlings. * and **: statistically significant difference compared to Col-0, one-tailed t-Test, 5% and 1% significance, respectively.

We next tested the function of *SPL9* in auxin responsiveness. To that end, we analyzed 3-day-old dark-grown Col-0, rSPL9, and *spl9-4* seedlings treated with ACC and the auxin inhibitor Yucasin^43^, which increases and decreases auxin levels in the inner cells of the apical hook, respectively^10^. The ethylene-signaling mutant *ein2-1* and the auxin-signaling mutant *arf3-2* were used as controls (Fig. 3b, c). We then measured the relative response (the fold-change between ACC and mock or between Yucasin and mock treatments) of the hook angle to the respective treatments. As expected, *ein2-5* exhibited reduced hook curvature and it was insensitive to ACC treatment^10^. Exogenous application of 10 µM of ACC increased hook angle similarly in rSPL9, *spl9-4,* and Col-0 (Fig. 3b), indicating that the miR156/*SPL9* module acts independently of ethylene to control auxin responses. Thus, the reduced expression of *MIR156C-GUS* at the hook region (Fig. 3a) was likely an indirect result of the ethylene-dependent increase of endogenous auxin in the hook. Accordingly, dark-grown rSPL9 seedlings were less sensitive to Yucasin treatment when compared to *spl9-4* and Col-0, which showed similar relative responses (Fig. 3c). The *arf3-2* mutant was insensitive to Yucasin and still able to establish the apical hook in the presence of the auxin-biosynthesis inhibitor (Fig. 3c). Collectively, our data suggest that the de-repression of *SPL9* is sufficient to bypass the normal requirement of high levels of endogenous auxin to promote hook curvature.

We next examined local auxin response in transgenic plants harboring the synthetic auxin-responsive DR5 promoter driving *GUS* expression (*DR5pro::GUS*). As previously reported^18^, *GUS* signal was observed in the concave side of the apical hook, mostly in the formation phase, and barely in the maintenance or opening phases (Supplementary Fig. 5a). On the other hand, auxin response was strongly intensified in the inner part of the rSPL9/*DR5pro::GUS* hook in the maintenance phase (Fig. 3d), indicating a local auxin maxima promoted by the de-repression of *SPL9* or loss of miR156-based regulation. To determine the molecular role of the miR156-targeted *SPL9* in the auxin pathway, we initially quantified the transcripts of several relevant genes associated with auxin biosynthesis, signaling, and response that are dependent on miR156 in the hypocotyl^28^. The auxin biosynthesis genes *YUCCA8 (YUC8)* and *-9 (YUC9)* were similarly expressed at the apices of dark-grown Col-0 and *spl9-4* seedlings, indicating that they are not dependent on *SPL9* regulation (Supplementary Fig. 5b). This result is line with previous observation that the miR156/*SPL9* module has only a marginal importance in regulating auxin biosynthesis^28^. On the other hand, we confirmed that at least one member of *Aux/IAA* and *SAUR* families was upregulated in the apical region of dark-grown *spl9-4* seedlings (Supplementary Fig. 5b), indicating that *SPL9* is necessary for their repression. In fact, *IAA19* and *SAUR19* were strongly downregulated at the apex of dark-grown rSPL9 seedlings compared to Col-0 (Fig. 3e). These results support the conclusion that *SPL9* is a repressor of auxin signaling and response genes, such as, *IAA19* and *SAUR19* at the apical region.

The auxin-mediated repression of *SAUR19* expression contributes to its inhibitory effect on PM H+-ATPase activity and consequent cell expansion in the hook^12^. This conclusion is supported by the constitutive expression of *SAUR19* from the 35S promoter (SAUR19oe plants), which results in increased hypocotyl cell length^44^. However, how *SAUR19* expression is modulated by auxin remains unclear^12^. One possibility is via the miR156/*SPL9* module, as *SAUR19* is repressed in MIM156 hypocotyl in response to warm temperature^33^ and in the rSPL9 apical region (Fig. 3e). To test whether miR156-targeted *SPL9*-based regulation of apical hook maintenance depends on *SAUR*, we generated rSPL9/SAUR19oe plants and used them in the time-lapse analysis of light triggering apical hook opening. As reported, *Arabidopsis* seedlings overexpressing *SAUR19* exhibited an accelerated apical hook opening. On the other hand, the presence of the *rSPL9* allele rescued the hook defects of SAUR19oe seedlings at wild-type levels (Fig. 4a). Thus, rSPL9/SAUR19oe plants provide genetic evidence supporting that *SPL9*-based regulation of apical hook maintenance is partially dependent on *SAUR19*.

Another possible downstream factor operating in the auxin-based regulation of cell growth in the early stages of etiolated seedling development is *FUL*, a hook opening repressor that suppresses *SAUR* expression^13,45^, and it was also activated in dark-grown rSPL9 apices, while repressed in *spl9-4* (Fig. 4b). We then generated rSPL9*/ful-7* seedlings and examined its apical hook opening response to light. While *ful-7* seedlings displayed an accelerated apical hook opening^37^, rSPL9*/ful-7* exhibited hook opening comparable to Col-0 (Fig. 4c). Thus, similar to *SAUR19*, *SPL9*-based regulation of apical hook maintenance is partially dependent on *FUL* function. We next asked whether SPL9-mediated *SAUR19* expression is dependent on *FUL*. To that end, we quantified *SAUR19* transcripts in apices of dark-grown *ful-7* and rSPL9*/ful-7* seedlings compared to Col-0. In line with the apical hook phenotype (Fig. 4a, c), *SAUR19* transcript levels at rSPL9/*ful-7* apices were comparable to that of Col-0 (Fig 4d). Collectively, our data indicate that miR156-targeted *SPL9* may be modulating auxin response by regulating *SAUR19* expression in both a *FUL*-dependent and independent way.

**Fig 4.**
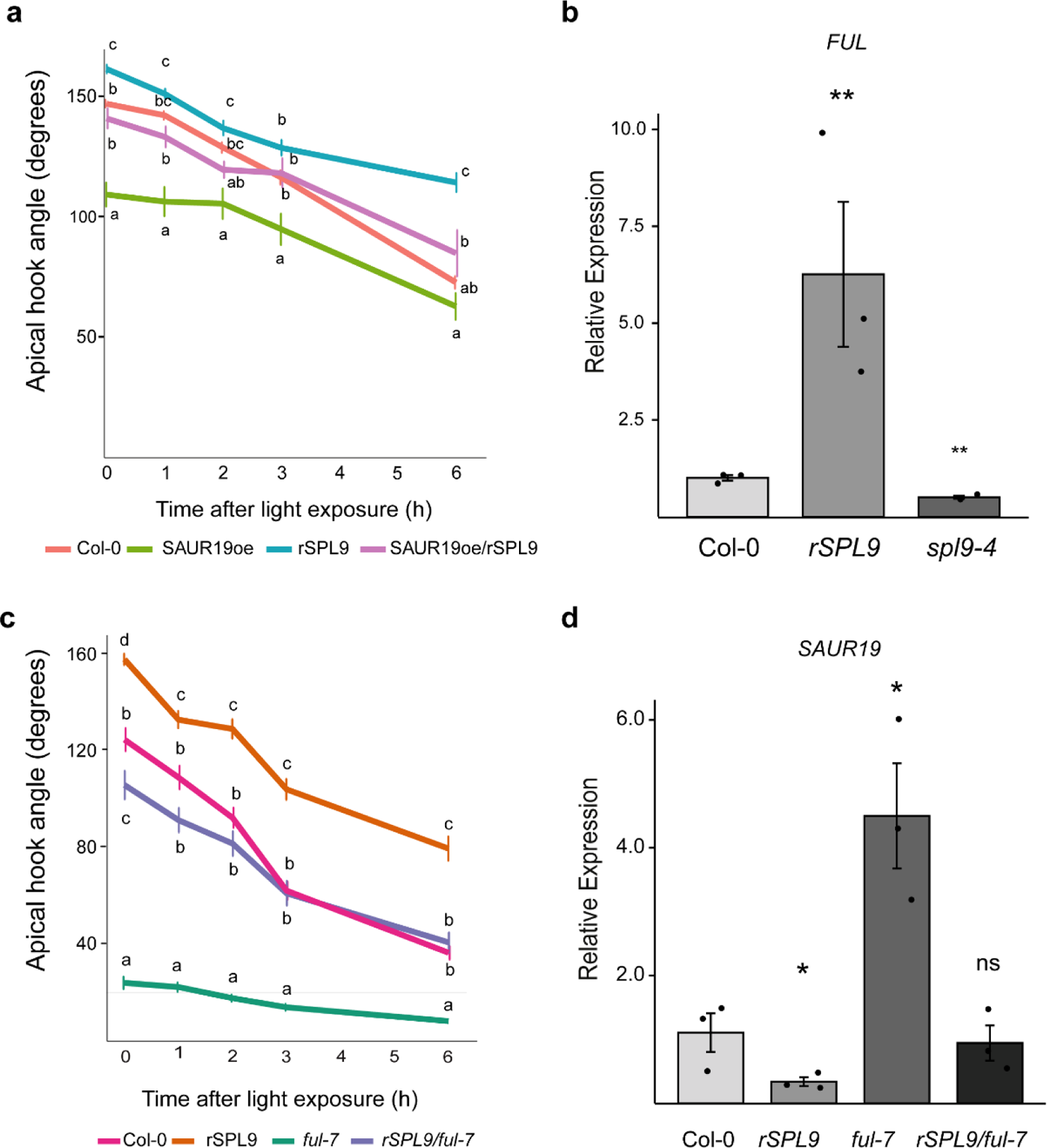
*SPL9*-based regulation of apical hook development is dependent on *SAUR19* and *FRUITFUL*. **a** Angle of the apical hook of 3-day-old dark-grown seedlings exposed to white light for 1, 2, 3, and 6 hours. Wild type (Col-0), *SAUR19* overexpression (SAUR19oe); miR156-resistant version of *SPL9* (rSPL9); and the double transgenic rSPL9/SAUR19oe. Different letters mean statistical significance among the genotypes at the same time point. ANOVA followed by Tukey test, 5% significance. **b** Relative expression of *FRUITFUL* (*FUL*) in Col-0, rSPL9, and the loss of function of *SPL9* (*spl9-4*). **: statistically significant difference from Col-0 by t-Test, p<0.01. **c** Angle of the apical hook of 3-day-old dark-grown seedlings exposed to white light for 1, 2, 3, and 6 hours. Col-0; rSPL9; loss of function of *FUL* (*ful-7*); and double transgenic-mutant rSPL9/*ful-7*. Different letters mean statistical significance among the genotypes at the same time point. ANOVA followed by Tukey test, 5% significance. **d** Relative expression of *SAUR19* in Col-0, rSPL9, *ful-7,* and rSPL9/*ful-7.* *: statistically significant difference from Col-0 by t-Test, p<0.05; ns: no statically significant difference.

*FUL* was already shown to be a direct target of SPLs^23^. To determine if *SAUR19* is a direct target of SPL9, we took advantage of the inducible expression system based on GR (Fig. 2c). rSPL9-GR seeds were plated on MS medium under dark conditions, and then treated with DEX in the presence or absence of cycloheximide (CHX) for 4 hours. CHX is a protein synthesis inhibitor, thus it blocks the translation of mRNAs regulated by SPL9, preventing secondary effects. We then quantified *SAUR19* transcripts at the apical region of dark-grown seedlings via qRT-PCR. We observed a similar reduction in *SAUR19* transcript levels (∼2-fold) in apices treated with DEX and with DEX+CHX (Fig. 5a), suggesting that *SAUR19* is a direct transcriptional target of SPL9. To further test whether SPL9 directly regulates *SAUR19* and *FUL*, we screened for putative SPL/SBP binding sites (bs) in both promoter regions. We found one and five SPL/SBP bs in *SAUR19* and *FUL* promoters, respectively. We then performed LUCIFERASE (LUC) transactivation assays by co-expressing *LUC* driven by either the *SAUR19* (*SAUR19pro::LUC*) or *FUL* promoter (upstream region plus 5’-UTR and first exon) (*FULpro::LUC*), with the *35Spro::rSPL9* or *35Spro::NLS-GFP* (negative control) constructs in *Nicotiana benthamiana* (*N. benthamiana*) leaves. The LUC activity of the *SAUR19pro::LUC* construct was significantly reduced in the presence of rSPL9 (Fig. 5b). On the other hand, rSPL9 was effective in activating *LUC* in the *FULpro::LUC* construct (Fig. 5c). Complementary DNA binding tests are needed to support the direct interaction of SPL9 and the *SAUR19* or *FUL* promoter regions, although the molecular and genetic results presented here strongly indicate an integrative action of miR156-*SPL9/SAUR19/FUL* during apical hook development.

**Fig 5.**
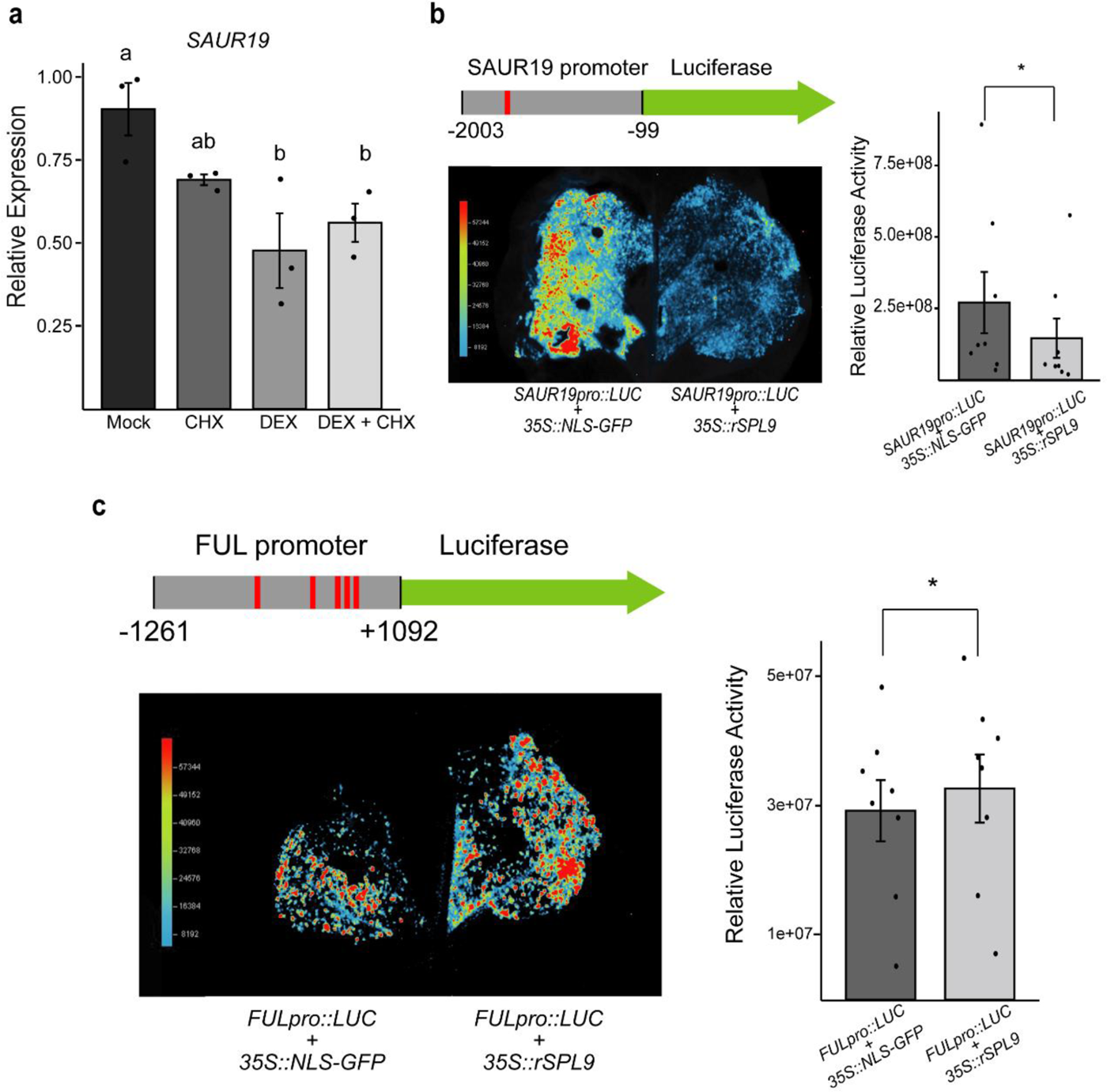
SPL9 directly regulates *SAUR19* and *FUL*. **a** Relative expression of *SAUR19* in dark-grown 3-d-old rSPL9-GR seedlings treated with Dexamethasone (DEX) in the absence or presence of cycloheximide (CHX). rSPL9-GR represses the expression of *SAUR19* in the absence of protein synthesis**. b-c.** Schematic representation of the 1904-bp *SAUR19* promoter region **(b)** or 2353-bp *FUL* promoter region **(c)** (gray rectangle) fused to *LUCIFERASE* (green arrow). The cis-element were identified as putative SPL/SBP binding sites (GTAC, red line). *Nicotiana benthamiana* leaves were co-infiltrated with Agrobacterium containing *35S::rSPL9* or *35S::NLS-GFP* (control) with *SAUR19pro::LUC* or *FULpro::LUC* constructs. Pseudo colors show the luminescence intensity (left). Quantification of relative luciferase activity (right) in leaves using NEWTON 7.0 imaging software. *: statistically significant difference by one-tailed t-Test, p<0.05.

### The miR156-dependent *SPL* regulation in skotomorphogenesis is critical for tomato seedling emergence

The role of *Arabidopsis SAUR19* in inhibiting PP2C.D during hypocotyl elongation is conserved in tomato seedlings^46^, and the overexpression of a miR156-resistant version of the *SPL9/15* tomato homolog *SlSBP15* reduces plant stature^30^. We then asked whether the miR156/*SPL* module is also important to control tomato seedling emergence. We evaluated seedling emergence rate at 3 days after germination of tomato cv. Micro-Tom (MT) and MT seedlings overexpressing a miR156-resistant version of *SlSBP15* (rSBP15), and the *CRISPR*-based knockout of *SlSBP15* (*sbp15*)^30,47^ (Fig. 6). Germinated seedlings from each genotype (Fig. 6a) were transferred to soil and covered with 0.5 cm of soil layer. High levels of *SlSBP15* in rSBP15 significantly repressed seedling emergence rate compared to MT and *sbp15* (Fig. 6b-d), reducing the stand for this genotype (Fig. 6c). To understand whether the miR156-targeted *SlSBP15* regulates tomato skotomorphogenesis, we measured the hypocotyl length of emerged seedlings 3 days after germination (Fig. 6e). De-repression of *SlSBP15* reduced tomato hypocotyl similar to what was observed for rSPL9 in *Arabidopsis* (Fig. 6e). Interestingly, the *sbp15* mutant showed an intermediary emergence rate and hypocotyl phenotype compared to MT and rSBP15 seedlings (Fig. 6d, e). The apparent reduction in hypocotyl length of both rSBP15 and *sbp15* seedlings is in line with the similar plant height phenotypes previously described^30^. Our results suggest that SPL/SBP proteins have a conserved role in *Arabidopsis* and tomato skotomorphogenesis, regulating cell expansion and impacting seedling emergence.

**Fig. 6.**
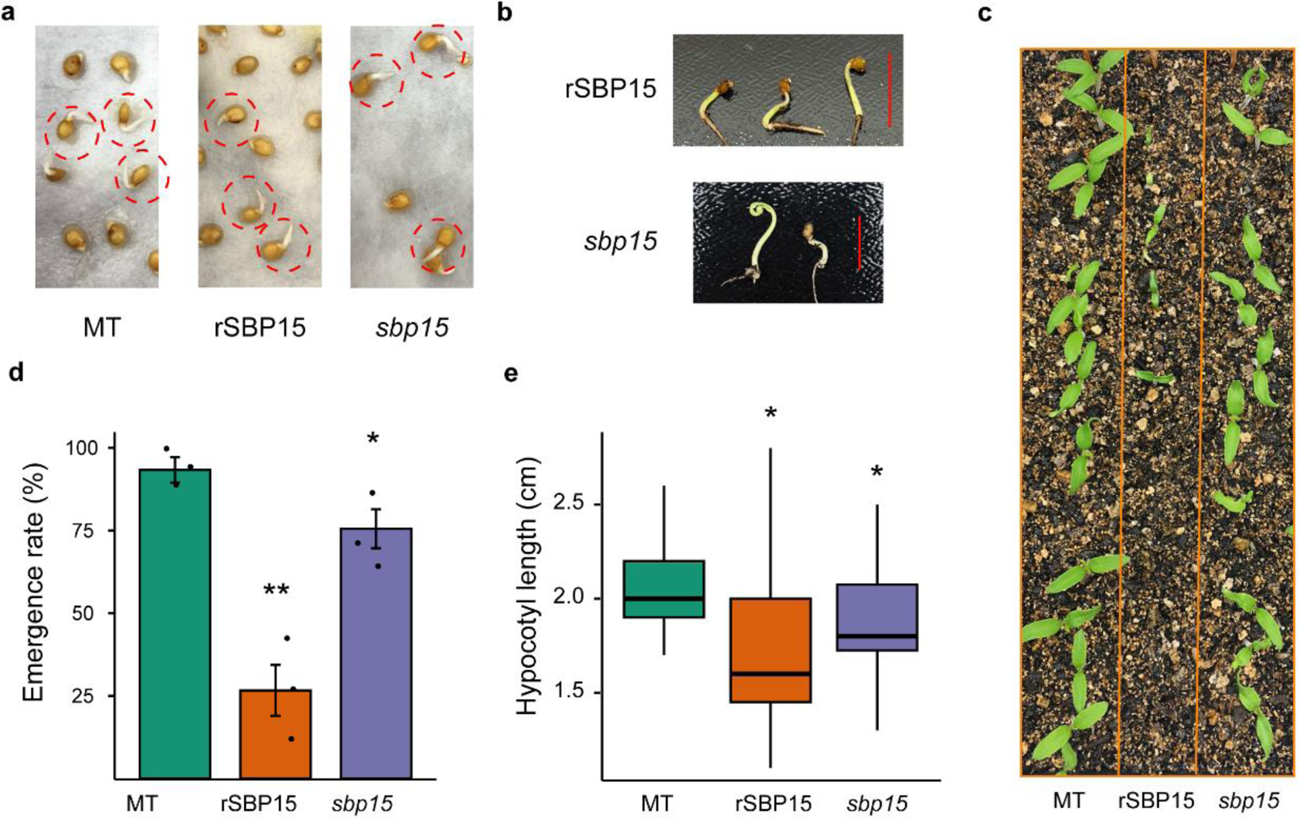
Tomato seedling emergence is affected by the de-repression of *SlSBP15*. **a** Germinated seedling phenotype (red circles) of wild type (MT), miR156 resistant version of *SlSBP15* (rSPB15); and *CRISPR*-based knockout of *SlSBP15* (*sbp15*). **b** non-emerged seedling phenotype of MT, rSPB15; and *sbp15*, 3 days after germination. Scale bar: 1 cm. **c** Emerged seedling phenotypes MT, rSPB15, and *sbp15*, 3 days after germination. **d** Percentage of emerged seedlings of MT, rSPB15, and *sbp15*, 3 days after germination in three replicates. * and **: Statistically difference from MT by t-Test, 5% and 1% significance, respectively. **e** Hypocotyl length of MT, rSPB15, and *sbp15*, 3 days after germination in three replicates. * and **: Statistically difference from MT by t-Test, 5% and 1% significance, respectively.

## DISCUSSION

When seeds germinate in the dark, skotomorphogenesis is activated in the seedling, promoting larger hypocotyls and the formation of the apical hook to favor emergence. Although it is well known that light and phytohormones, including ethylene and auxin, modulate apical hook formation ^35,48,49^, how miRNA-controlled modules participate in this process remains uncertain. Here, we showed that miR156-based regulation of *SPL9* in the inner part of the apical hook is sufficient and necessary for the proper coordination of cell growth. Although it has been postulated that mutations in core enzymes of the miRNA biogenesis, such as SE, affect skotomorphogenesis^18,50^, the link between specific miRNAs and differential cell expansion during apical hook formation has not been previously reported. In addition, whether and how the miR156 module mediates auxin-dependent *SAUR19* repression during hook maintenance was previously unknown. Thus, the miR156/*SPL9* module defines an unanticipated genetic circuit that connects asymmetric auxin distribution and the repression of *SAUR19* activity to prevent the apical hook from opening in the maintenance phase (Fig. 7a).

**Fig. 7.**
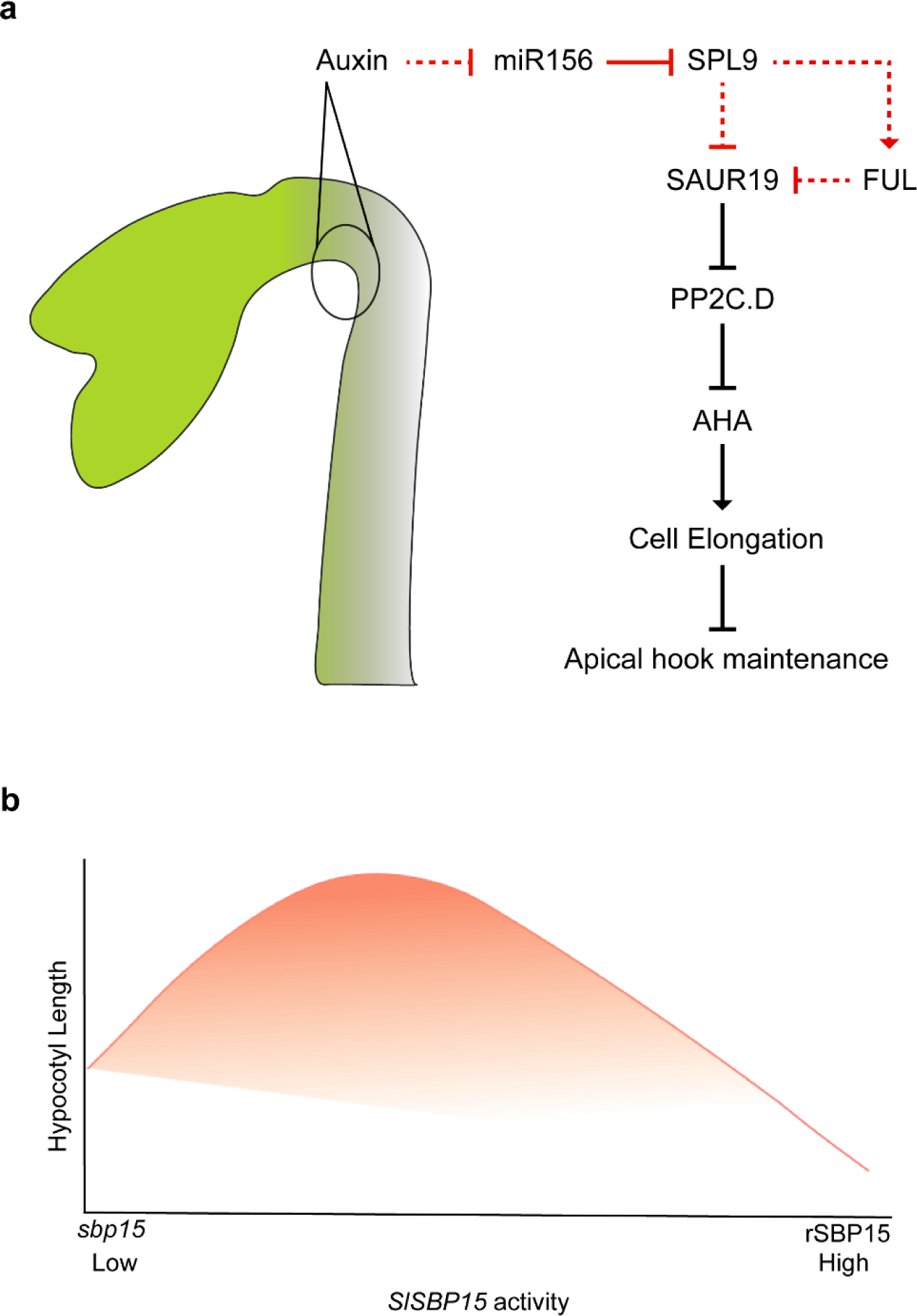
Proposed model of the genetic circuity including Auxin/miR156-*SPL9*/*FUL*/*SAUR19* in *Arabidopsis* apical hook and the relationship between *SlSBP15* expression levels and tomato hypocotyl growth. **a** The miR156/SPL module acts as a hub between the auxin and the *FUL/SAUR* pathway to regulate cell expansion mediated by the autoinhibited H+-ATPase domain (AHA) during the apical hook development. **b** Representation that lower or higher levels of *SBP15* transcripts in tomato seedlings lead to shorter hypocotyls.

Both apical hook and hypocotyl development are coordinated mainly by cell expansion^5,51^. Our results show that the miR156-dependent regulation is required for both hook and hypocotyl development elicited by modified cell growth under dark conditions. Genetic evidence supports the conclusion that altered levels of miR156 dramatically affected hypocotyl elongation and hook opening in dark-grown seedlings (Fig. 1), which is in line with recent reports that SE and DCL1 functions are crucial to promote proper skotomorphogenetic responses^17,50^. Although it was not the main focus of this study, our data also suggest that miR156 regulation is important to control hypocotyl growth in diurnal conditions, as MIM156 and *mir156* quadruple mutant exhibited shorter hypocotyls under dark conditions, and also after 6h of white light treatment (Supplementary Fig.1). Among the miR156-targeted *SPLs*, we show here that the de-repression of *SPL9* appears to have the most prominent effect on hook formation (Fig. 1). In fact, stabilizing *SPL9* RNA alone in the rSPL9 line, mainly during skotomorphogenesis, is sufficient to promote rSPL9 protein accumulation in the inner part of apical hook (Fig. 1g, h), and to repress cell growth (Fig. 2). *SPL9* is part of the group of *SPLs* related to the control of phase transition^40^. At the seedling stage, a remarkable transition from skotomorphogenesis to photomorphogenesis is induced through environmental cues and differential gene expression profiles^52^. Analyzing publicly available high throughput degradome data of de-etiolated *Arabidopsis* seedlings^20^, we noticed that *SPL9* displayed the highest ratio of light to dark degradome signature among the miR156-targeted *SPLs*, followed by its homologous *SPL15* (Supplementary Table 1). This observation, along with data showing that the miR156-based regulation by *SPL9* in the early stages of seedling development is pivotal to modulating hook opening in response to light (Fig. 2c and Supplementary Fig. 4), provides strong evidence that miR156 enables the dark to light growth plasticity of *Arabidopsis* seedlings by repressing mainly *SPL9*. At this point, however, we cannot rule out the possibility that other *SPLs* may directly participate in miR156-dependent hook opening in response to light. For instance, the de-repression of *SPL15* led to a higher hook angle, which was comparable to that of rSPL9 seedlings under dark conditions (Fig. 1).

Asymmetric auxin gradient is essential to the apical hook establishment, and plants that are either overproduced or are auxin-resistant are hookless^48,53^. We observed that the exogenous application of Picloram, which likely disrupted the endogenous auxin gradient, reduced *MIR156C-GUS* expression at the apical hook region (Fig. 3). This indicates that auxin may regulate *SPL9* expression in a miR156-dependent manner at the hook under dark conditions (Fig. 7a). This conclusion is supported by the fact that dark-grown rSPL9 seedlings were less sensitive to Yucasin, and *spl9-4* seedlings were as sensitive to the auxin inhibitor as Col-0 (Fig. 3b). Because rSPL9 were similarly sensitive to exogenous application of ACC as Col-0 seedlings, and despite the fact that high ACC levels repressed *MIR156C-GUS* (Fig. 3), it is unlikely that miR156 is a major contributor to the auxin-ethylene interactions during apical hook formation. Nonetheless, miR156 and its targets were documented to be essential to control ethylene production during blueberry and tomato fruit ripening^42,54^, suggesting a developmental-dependent direct interplay between miR156 and ethylene. Previous reports indicated that the miR156/*SPL* “hub” affects auxin transport during light-associated developmental contexts^25,30,47^, and in this study, we demonstrated how miR156 is essential to mediate auxin responsiveness during skotomorphogenesis.

The miR156/*SPL* module has been associated with the control of cell number and size in meristematic regions and leaves^47,55,56^. Our results revealed that miR156-targeted *SPL9* has an essential role in enabling cell growth through the modulation of auxin-responsive genes. Auxin-responsive SAUR19 binds to PP2C-D phosphatases inhibiting their activity on the PM H^+^-ATPase, and consequently promoting cell expansion^44^. In contrast to rSPL9, dark-grown *SAUR19*-overexpressing seedlings exhibit larger hypocotyls and more open apical hook^57^. Here, we propose that miR156-targeted *SPL9* is one of the main contributors to the auxin-dependent *SAUR19* regulation (Fig. 7a). This conclusion is supported by several lines of evidence: (1) *SAUR19* is downregulated in the concave side of the apical hook in response to higher auxin concentration, but this mechanism does not depend directly on *ARF7*^12^; (2) Our transcript quantification and luciferase assays supported the idea of more direct repression of *SAUR19* by SPL9 (Fig. 3e, 5a); (3) The rSPL9/SAUR19oe line rescued the wild type Col-0 hook phenotype, indicating that a proper activity of *SAUR19* is at least partially necessary for the rSPL9 hook phenotype (Fig. 4a); (4) In response to elevated temperature, miR156 conditions auxin sensitivity to promote hypocotyl elongation by regulating members of *SAUR* family^28^. In addition, SPL9 may regulate *SAUR19* through the activation of *FUL*. FUL is reported as an apical hook opening repressor that acts by suppressing *SAUR* expression^13,45^. During *Arabidopsis* flowering, *FUL* is targeted by SPLs^31^. We showed here that this regulation may be conserved in the apical hook (Fig. 7a). *FUL* was upregulated in the apices of rSPL9 seedlings (Fig. 4b), and the *FUL* promoter was activated by rSPL9 in *N. benthamiana* leaves (Fig 5b). More importantly, the rSPL9/*ful-7* line also rescued hook phenotype and the *SAUR19* transcript levels at the wild type levels (Fig 5c, d). Collectively, these findings indicate that the miR156/*SPL9* module controls auxin-mediated hook formation and maintenance by repressing *SAUR19* (and likely other auxin signaling genes) in a *FUL*-dependent and independent manner (Fig. 7a).

From an agronomical point of view, understanding the skotomorphogenetic seedling development may enable making better decisions regarding sowing depth and seed number, which are reflected in stand establishment and yield^2,58^. When the apical hook opens too early, the shoot apical meristem and cotyledons can be damaged, affecting the seedling survival. For instance, *SAUR19* overexpression in *Arabidopsis* or tomato led to a more open apical hook (Fig. 5a), and reduced emergence rate^12^. Similarly, shorter hypocotyls may also indicate a delay in seedling growth resulting in unsuccessful emergence. In tomato, an important worldwide crop, high levels of a miR156-resistant *SlSBP15* version^30,47^ in rSBP15 seedlings led to shorter hypocotyl and reduced seedling emergence rate (Fig 6). Strikingly, the knockout of *SlSBP15* led to an intermediate hypocotyl length between that of MT and rSBP15 affecting the emergence rate (Fig 6d, e). Altogether, these findings suggest that miR156 plays an essential role in enabling *SlSBP15* function during early seedling growth and that a fine-tuned regulation of *SPL/SBP* activity during hypocotyl and apical hook growth may improve seedling vigor and emergence (Fig. 7b).

In summary, our results indicate a new role of the miR156/*SPL9* module in the control of seedling emergence (Fig. 7). We propose the existence of the circuity Auxin/miR156-*SPL9/FUL/SAUR* as a molecular link between auxin maxima and *SAUR19* activity in the inner part of the apical hook. Our study contributes to elucidate the mechanism by which miR156 and its targets mediate auxin responses in the hook, and may provide new targets to improve seedling emergence vigor.

## METHODS

### Plant material and growth conditions

All *Arabidopsis* (*Arabidopsis thaliana*) lines used in this study are in Columbia (Col-0) background. The mutants and transgenic lines: miR156oe (*pro35S::MIR156A*)^34^; MIM156 (*pro35S::MIM156*)^34^; *MIR156A-GUS* (*proMIR156A::MIR156A-GUS*)^38^; *MIR156C-GUS* (*proMIR156C::MIR156C-GUS*)^38^; GFP sensors control (*proUBQ10:GFP-N7-TNOS*) and GFP-miR156 (*proUBQ10:GFP-N7-SPL3 3’UTR*)^39^; *rSPL3-GUS* (*proSPL3::rSPL3-GUS*)^40^; *rSPPL6-GUS* (*proSPL6::rSPL6-GUS*)^40^; *rSPL9-GUS* (*proSPL9::rSPL9-GUS*)^40^; *rSPL10-GUS* (*proSPL10::rSPL10-GUS*)^40^; *sSPL9-GUS* (*proSPL9::sSPL9-GUS*)^40^; rSPL9 (*proSPL9::rSPL9*)^59^; rSPL9-GR (*proSPL9::rSPL9-GR*)^23^; GFP-rSPL9 (*proSPL9::GFP-rSPL9*)^59^; *spl9/15* (*spl9-4 spl15-1*)^59^; *ein2-5*^9^; *arf3-2*^60^; DR5pro::GUS^61^; *ful-7*^45^; SAUR19oe (*35S:StrepII-SAUR19*)^57^; were previously described. The mutants *mir156a/b/c/d* (CS71704); *spl9-4* (CS807258); *q-spl* (*spl2/9/11/13/15*, CS69798); and rSPL15 (CS2109787) were obtained from the *Arabidopsis* Biological Resource Center. Double transgenic/mutant DR5::GUS/rSPL9; rSPL9/*ful-7* were obtained by crossing and selection of the homozygous in F3 generation. Double transgenic rSPL9/SAUR19oe were obtained by crossing and analyzed in F1 generation. Tomato (*Solanum lycopersicum*) lines rSBP15 and *sbp15* were in Micro-Tom LA3911 (MT) background and were previously described ^30^.

*Arabidopsis* seeds were sterilized in 70% ethanol with tween for 10 minutes and washed in sterile water three times. The seeds were plated on half-strength Murashige and Skoog (½ MS) basal medium with Gamborg’s vitamins (Sigma Aldrich, M0404), pH 5.7, and 0.8% agar. The seeds were stratified at 4°C for 48 hours in darkness, then transferred to white light for four hours to stimulate germination, and subsequently covered with two layers of aluminum foil and left vertically at 22°C for the 3 days. For crossings, seeds were stratified in 0.1% agarose solution at 4°C for 48 hours in darkness and individually sown in peat soil SPHAGNUM TPS JIFFY® with vermiculite (2:1) and cultivated at 22°C, 16 h photoperiod. Crossings were performed as indicated^62^.

### Hypocotyl length and apical hook angle measurements

For the hypocotyl and apical hook measurements, at least 40 seedlings from each genotype were scanned using an Epson V370 photo scanner. The hypocotyl (distance between the cotyledon and the roots) and the apical hook (complementary angle between the hypocotyl axes and the cotyledon) were measured using NIH ImageJ software (http://rsb.info.nih.gov/nih-image/). For light treatment, the dark-grown seedlings were scanned and left in white light, followed by scanning after 1, 2, 3, and 6 hours. The relative phytohormone response in the apical hook was calculated by dividing the angle of treated seedlings by the respective Mock, three replicates were used to calculate mean and standard error. Graphs were plotted using ggplot2 in R software. Statistical analyses were performed in InfoStat-Statistical Software. Representative pictures were taken in a Nikon SMZ18 stereomicroscope with a zoom of 6x and processed in Adobe Illustrator software.

### Vector constructs and plant transformation

To obtain transgenic plants expressing *SPL9pro::GUS* three fragments of the intergenic region (2kb) upstream to *SPL9* were individually cloned into pUPD2 vector and then ligated upstream to the *β-glucuronidase (GUS)* (pUPD2) and *35S* terminator (pLOM) into the pICSL86900_OD vector by Golden Gate cloning^63,64^. The *proSPL9::GUS* construction was agroinfiltrated in Col-0 plants by the floral dip method as previously described^65^. Several independent lines were obtained, and three were further analyzed.

For transactivation assays, two fragments of *SAUR19* promoter and *FUL* promoter with the 5’ UTR and first exon were individually cloned into pUPD2. *SAUR19* and *FUL* fragments were then ligated upstream of the firefly *LUCIFERASE* (*LUC*) and *35S* terminator in the pLOM vector^30^. To obtain a miR156-resistant version of *SPL9* (rSPL9) synonymous mutations into the miR156-binding site was introduced by PCR. Two fragments of rSPL9, with domesticated Esp31 restriction site, were cloned into pUPD2 and then ligated downstream to 35S promoter (pUPD2) into the pICSL86900_OD to generate the 35S::rSPL9 construct. Oligonucleotide sequences are listed in Supplementary Table 2.

### Histochemical β-glucuronidase (GUS) staining

For the expression analysis via GUS staining the seedlings were harvested and fixed in 90% acetone in the darkroom with a green light source. The GUS staining protocol was performed as described^62^ and incubated from 45 min to overnight at 37°C. For each genotype/treatment were analyzed at least 10 seedlings. Representative images were taken in a Nikon SMZ18 stereomicroscope with zoom of 13.5x and processed in Adobe Photoshop CS6 and Adobe Illustrator software.

### Phytohormones and chemical treatments

The stock solutions were prepared using DMSO for ACC (10 mM), Picloram (5 mM), Yucasin (50 mM), Dexamethasone (10 mM), and cycloheximide (10 mM). Seeds were plated on MS medium supplemented with Mock (0.1% DMSO), ACC (10 μM), Picloram (5 μM), and Yucasin (50 μM) and grown in the dark for 3 days. For Dexamethasone treatment, seeds were plated on MS medium supplemented with Mock (0.1% DMSO) or Dexamethasone (10 μM) and grown in the dark. After three days, the plates were opened in a sterile hood in the darkroom under green light and received 1 mL of either Mock (0.1% DMSO) or DEX (10 μM). The plates were incubated in the growth chamber under white light for 3 hours and scanned. For DEX and cycloheximide (CHX) expression analyses, rSPL9-GR seedlings were grown in 1/2X MS medium in the dark. After three days, the plates were opened in a sterile hood in the darkroom under green light and received 1 mL of Mock (0.1% DMSO), CHX (10 μM), DEX (10 μM), or DEX + CHX (10 μM of each) in three replicates and kept in the dark for 4 hours. The apices of the seedlings were dissected and harvested, frozen in liquid nitrogen, and qRT-PCR was performed as described bellow.

### Fluorescence microscope and confocal images

For the GFP sensor assay, 3-day-old dark-grown seedlings were kept in dark or exposed to white light for 3 hours and immediately imaged in a fluorescence microscope Zeiss Axio Imager.A2 ate 20x objective. Images from at least ten seedlings from each genotype were taken and processed using Photoshop CS6 Software. For cell size analyses, 3-day-old dark-grown seedlings from Col-0, rSPL9, and *spl9-4* were previously incubated in a solution of 10 μg of propidium iodide for 10 minutes, rinsed in water and imaged in confocal microscope Nikon C2+, excitation laser 405 nm. Cell size was measured using ImageJ software. Graphs were plotted using ggplot2 in R, and ANOVA followed by Tukey test was performed in InfoStat-Statistical Software. For GFP-rSPL9 images, 3-day-old dark-grown seedlings were incubated in propidium iodide as above, and imaged in a confocal microscope Nikon C2+, excitation lasers 405 nm (propidium iodide) and 488 nm (GFP). Images were processed in Adobe Photoshop CS6 Software.

### RNA extraction and qRT-PCR analysis

The apical region of 3-day-old dark-grown seedlings, was dissected in the dark with a green light source and immediately frozen in liquid nitrogen. Total RNA was extracted by Trizol reagent (Invitrogen) protocol, followed by DNase I (Invitrogen) treatment. The complementary DNA (cDNA) synthesis was performed with 1 μg of total RNA using ImProm-II™ Reverse Transcriptase (Promega). cDNA was diluted in DNase-free water and used for qRT-PCR with Go Taq® qPCR Master Mix reagent (PROMEGA) and analyzed in a Step-OnePlus real-time PCR system (Applied Biosystems). For each genotype, at least three biological samples with three technical replicates were used in the qRT–PCR analyses. The *ACTIN2* gene expression was selected as an internal control. Oligonucleotide sequences are listed in the Supplementary Table 2. Graphs were plotted using ggplot2 in R software and two-tailed t-test analyses or ANOVA followed by Tukey’s test were performed.

### Luciferase transactivation assays

Each reporter construct (*SAUR19pro::LUC* or *FULpro::LUC*) combined to the effector (*35S::rSPL9*) or the negative control (*35S::NLS-GFP*^30^) were agroinfiltrated into 5-week-old *Nicotiana benthamiana* leaves. After 3 days the leaves were sprayed with D-lucifernin (25mg/L) (Promega) and LUC activity was evaluated using the NEWTON 7.0 CCD imagin system (Vilber). Relative LUC activity in leaves was quantified using the NEWTON 7.0 software (https://www.vilber.com/newton-7-0/) with default parameters. At least seven leaves were evaluated to calculate the mean and standard deviation. Graphs were plotted using ggplot2 in R software and two-tailed t-test analyses were made in Microsoft office Excel software.

### Tomato emergence rate

Tomato seeds of wild type (MT), rSBP15, and *sbp15* were sterilized in 5% sodium hypochlorite for five minutes and rinsed with distilled water three times, the seeds were disposed in individual dark germination box containing four layers of humid filter papers. After two days, 15 germinated seeds from each genotype were transferred to soil in 5 mm depth in three replicates. The number of seedlings that emerged was counted at three days after seeds transfer. Emerged seedlings were taken from the soil and the hypocotyl length was measured using a pachymeter. The non-emerged seedlings from rSBP15 and *sbp15* lines were rescued from the soil and photographed. The percentage of emerged seedlings was calculated. The graphs were plotted using ggplot2 in R software. ANOVA and Tukey tests were performed in InfoStat-Statistical Software.

### Accession numbers

*MIR156A* (AT2G25095), *MIR156B* (AT4G30972)*, MIR156C* (AT4G31877)*, MIR156D* (AT5G10945)*, SPL2* (AT5G43270), *SPL3* (AT2G33810)*, SPL6* (AT1G69170), *SPL9* (AT2G42200), *SPL10* (AT1G27370), *SPL11* (AT1G27360), *SPL13* (AT5G50570), *SPL15* (AT3G57920), *EIN2* (AT5G03280), *ARF3* (AT2G33860), *FUL* (AT5G60910), *SAUR 10* (AT2G18010), *SAUR19* (AT5G18010), *SAUR26* (AT3G03850), *YUC8* (AT4G28720), *YUC9* (AT1G04180), *IAA19* (AT3G15540), *IAA29* (AT4G32280), *ACTINA2* (AT3G18780), SlSBP15 (Solyc10g078700).

## Acknowledgments

We thank the members of Dr Nogueira and Dr Pedmale laboratories for helpful discussions. We Thank Dr. William Gray (University of Minnesota), Dr. Jia-wei Wang (Center for Excellence in Molecular Plants Science), and Dr. Marian Bemer (Wageningen University & Research) for providing plant material.

## Data availability

All relevant data can be found within the article and its supplementary information, and they are openly available upon request.

## Author contributions

F.G.P and F.T.S.N. were responsible for the conception, planning and organization of the experimental time line. F. G. P.; A. J. M. S.; M. H. V.; and L. T. carried out the experiments. F.T.S.N. and U.P. directly supervised the experimental work done by F. G. P.; A. J. M. S.; M. H. V.; and L. T. F.G.P and F.T.S.N. wrote the manuscript, and U.P. helped to revise the manuscript.

## REFERENCES

1. Nemhauser, J. & Chory, J. Photomorphogenesis. Arabidopsis Book 1, e0054 (2002).

2. Samiei, S., Rasti, P., Ly Vu, J., Buitink, J. & Rousseau, D. Deep learning-based detection of seedling development. Plant Methods 16, (2020).

3. Josse, E. M. & Halliday, K. J. Skotomorphogenesis: The Dark Side of Light Signalling. Current Biology 18, 1144–1146 (2008).

4. Darwin, C. & Darwin, F. *The Power of Movement in Plants.* (D. Appleton and Co, New York, 1881).

5. Raz, V. & Ecker, J. R. Regulation of differential growth in the apical hook of Arabidopisis. Development 126, 3661–3668 (1999).

6. Liscum, E. & Hangarter, R. P. Photomorphogenic mutants of Arabidopsis thaliana reveal activities of multiple photosensory systems during light-stimulated apical-hook opening. Planta 191, (1993).

7. Wang, Y., Peng, Y. & Guo, H. To curve for survival: Apical hook development. J Integr Plant Biol 65, 324–342 (2023).

8. Rolfe, B. et al. The Ethylene Signal Transduction Pathway in Plants. Science (1979) 268, 667–675 (1995).

9. Guzmán, P. & Ecker, J. R. Exploiting the triple response of Arabidopsis to identify ethylene-related mutants. Plant Cell 2, 513–523 (1990).

10. Vandenbussche, F. et al. The auxin influx carriers AUX1 and LAX3 are involved in auxin-ethylene interactions during apical hook development in Arabidopsis thaliana seedlings. Development 137, 597–606 (2010).

11. Abbas, M., Alabadí, D., Blázquez, M. A., Pollmann, S. & Elio, F. Differential growth at the apical hook: all roads lead to auxin. Front Plant Sci 4, (2013).

12. Du, M. et al. Biphasic control of cell expansion by auxin coordinates etiolated seedling development. Sci Adv 8, 1570 (2022).

13. Führer, M. et al. FRUITFULL is a Repressor of Apical Hook Opening in Arabidopsis thaliana. International Journal of Molecular Sciences 2020, Vol. 21, Page 6438 21, 6438 (2020).

14. Lin, M. C., Tsai, H. L., Lim, S. L., Jeng, S. T. & Wu, S. H. Unraveling multifaceted contributions of small regulatory RNAs to photomorphogenic development in Arabidopsis. BMC Genomics 18, (2017).

15. Tsai, H. L. et al. HUA ENHANCER1 is involved in posttranscriptional regulation of positive and negative regulators in Arabidopsis photomorphogenesis. Plant Cell 26, 2858–2872 (2014).

16. Voinnet, O. Leading Edge Review Origin, Biogenesis, and Activity of Plant MicroRNAs. Cell 136, 669–687 (2009).

17. Sacnun, J. M., Crespo, R., Palatnik, J., Rasia, R. & González-Schain, N. Dual function of HYPONASTIC LEAVES 1 during early skotomorphogenic growth in Arabidopsis. Plant Journal 102, 977–991 (2020).

18. Vacs, P., Rasia, R. & González-Schain, N. HYPONASTIC LEAVES 1 is required for proper establishment of auxin gradient in apical hooks. Plant Physiol 187, 2356–2360 (2021).

19. Li, Y., Varala, K. & Hudson, M. E. A survey of the small RNA population during far-red light-induced apical hook opening. Front Plant Sci 5, (2014).

20. Lin, M. C., Tsai, H. L., Lim, S. L., Jeng, S. T. & Wu, S. H. Unraveling multifaceted contributions of small regulatory RNAs to photomorphogenic development in Arabidopsis. BMC Genomics 18, (2017).

21. Wu, G. & Poethig, R. S. Temporal regulation of shoot development in Arabidopsis thaliana by miRr156 and its target SPL3. Development 133, 3539–3547 (2006).

22. Morea, E. G. O. et al. Functional and evolutionary analyses of the miR156 and miR529 families in land plants. BMC Plant Biol 16, 40 (2016).

23. Wu, G. et al. The Sequential Action of miR156 and miR172 Regulates Developmental Timing in Arabidopsis. Cell 138, 750–759 (2009).

24. Silva, G. F. F. et al. Tomato floral induction and flower development are orchestrated by the interplay between gibberellin and two unrelated microRNA-controlled modules. New Phytologist 221, 1328–1344 (2019).

25. Barrera-Rojas, C. H. et al. miR156-targeted SPL10 controls root meristem activity and root-derived de novo shoot regeneration via cytokinin responses. J Exp Bot (2019) doi:10.1093/jxb/erz475.

26. Silva, G. F. F. E. et al. MicroRNA156-targeted SPL/SBP box transcription factors regulate tomato ovary and fruit development. Plant Journal 78, 604–618 (2014).

27. 27. Xie, Y., et al. Phytochrome-interacting factors directly suppress MIR156 expression to enhance shade-avoidance syndrome in Arabidopsis. Nat Commun 8, (2017).

28. Sang, Q. et al. MicroRNA156 conditions auxin sensitivity to enable growth plasticity in response to environmental changes in Arabidopsis. Nat Commun 14, (2023).

29. Zhao, J., Shi, M., Yu, J. & Guo, C. SPL9 mediates freezing tolerance by directly regulating the expression of CBF2 in Arabidopsis thaliana. BMC Plant Biol 22, (2022).

30. Barrera-Rojas, C. H. et al. Tomato miR156-targeted SlSBP15 represses shoot branching by modulating hormone dynamics and interacting with GOBLET and BRANCHED1b. J Exp Bot (2023) doi:10.1093/jxb/erad238/7205173.

31. Yamaguchi, A. et al. The MicroRNA-Regulated SBP-Box Transcription Factor SPL3 Is a Direct Upstream Activator of LEAFY, FRUITFULL, and APETALA1. Dev Cell 17, 268–278 (2009).

32. Wang, J.-W., Schwab, R., Czech, B., Mica, E. & Weigel, D. Dual Effects of miR156-Targeted SPL Genes and CYP78A5/KLUH on Plastochron Length and Organ Size in Arabidopsis thaliana. Plant Cell 20, 1231–1243 (2008).

33. Nodine, M. D. & Bartel, D. P. MicroRNAs prevent precocious gene expression and enable pattern formation during plant embryogenesis. Genes Dev 24, 2678– 2692 (2010).

34. Franco-Zorrilla, J. M. et al. Target mimicry provides a new mechanism for regulation of microRNA activity. Nat Genet 39, 1033–1037 (2007).

35. Liscum, E. & Hangarter, R. P. Light-Stimulated Apical Hook Opening in Wild-Type Arabidopsis thaliana Seedlings. Plant Physiol 101, 567–572 (1993).

36. Fouracre, J. P., He, J., Chen, V. J., Sidoli, S. & Poethig, R. S. VAL genes regulate vegetative phase change via miR156-dependent and independent mechanisms. PLoS Genet 17, e1009626 (2021).

37. He, J. et al. Threshold-dependent repression of SPL gene expression by miR156/miR157 controls vegetative phase change in Arabidopsis thaliana. PLoS Genet 14, e1007337 (2018).

38. Yang, L., Xu, M., Koo, Y., He, J. & Scott Poethig, R. Sugar promotes vegetative phase change in Arabidopsis thaliana by repressing the expression of MIR156A and MIR156C. Elife 2013, (2013).

39. Cheng, Y.-J. et al. Cell division in the shoot apical meristem is a trigger for miR156 decline and vegetative phase transition in Arabidopsis. PNAS 118, 3–9 (2021).

40. Xu, M. et al. Developmental Functions of miR156-Regulated SQUAMOSA PROMOTER BINDING PROTEIN-LIKE (SPL) Genes in Arabidopsis thaliana. PLoS Genet 12, 1–29 (2016).

41. Guo, A. Y. et al. Genome-wide identification and evolutionary analysis of the plant specific SBP-box transcription factor family. Gene 418, 1–8 (2008).

42. Li, H. et al. The miR156/SPL12 module orchestrates fruit colour change through directly regulating ethylene production pathway in blueberry. Plant Biotechnol J 22, 386–400 (2024).

43. Nishimura, T. et al. Yucasin is a potent inhibitor of YUCCA, a key enzyme in auxin biosynthesis. The Plant Journal 77, 352–366 (2014).

44. Spartz, A. K. et al. SAUR Inhibition of PP2C-D Phosphatases Activates Plasma Membrane H +-ATPases to Promote Cell Expansion in Arabidopsis C W. Plant Cell 26, 2129–2142 (2014).

45. Bemer, M. et al. FRUITFULL controls SAUR10 expression and regulates Arabidopsis growth and architecture. J Exp Bot 68, 3391–3403 (2017).

46. Spartz, A. K. et al. Constitutive Expression of Arabidopsis *SMALL AUXIN UP RNA19* (*SAUR19*) in Tomato Confers Auxin-Independent Hypocotyl Elongation. Plant Physiol 173, 1453–1462 (2017).

47. Ferigolo, L. F. et al. Gibberellin and miRNA156-targeted SlSBP genes synergistically regulate tomato floral meristem determinacy and ovary patterning. Development 150, (2023).

48. Žádníková, P., et al. Role of PIN-mediated auxin efflux in apical hook development of Arabidopsis thaliana. Development 137, 607–617 (2010).

49. Zhang, X. et al. Integrated regulation of apical hook development by transcriptional coupling of EIN3/EIL1 and PIFs in Arabidopsis. Plant Cell 30, 1971–1988 (2018).

50. Sun, Z. et al. Coordinated regulation of Arabidopsis microRNA biogenesis and red light signaling through Dicer-like 1 and phytochrome-interacting factor 4. PLoS Genet 14, e1007247 (2018).

51. Derbyshire, P., Findlay, K., Mccann, M. C. & Roberts, K. Cell elongation in Arabidopsis hypocotyls involves dynamic changes in cell wall thickness. J Exp Bot 58, 2079–2089 (2007).

52. Ma, L. et al. Light Control of Arabidopsis Development Entails Coordinated Regulation of Genome Expression and Cellular Pathways. Plant Cell 13, 2589– 2607 (2001).

53. Wang, Y. & Guo, H. On hormonal regulation of the dynamic apical hook development. New Phytologist 222, 1230–1234 (2019).

54. Adaskaveg, J. A., Silva, C. J., Huang, P. & Blanco-Ulate, B. Single and Double Mutations in Tomato Ripening Transcription Factors Have Distinct Effects on Fruit Development and Quality Traits. Front Plant Sci 12, (2021).

55. Fouracre, J. P. & Poethig, R. S. Role for the shoot apical meristem in the specification of juvenile leaf identity in *Arabidopsis*. Proceedings of the National Academy of Sciences 116, 10168–10177 (2019).

56. Usami, T., Horiguchi, G., Yano, S. & Tsukaya, H. The more and smaller cells mutants of Arabidopsis thaliana identify novel roles for SQUAMOSA PROMOTER BINDING PROTEIN-LIKE genes in the control of heteroblasty. Development 136, 955–964 (2009).

57. Spartz, A. K. et al. The SAUR19 subfamily of SMALL AUXIN UP RNA genes promote cell expansion. The Plant Journal 70, 978–990 (2012).

58. Kong, X. et al. Monoseeding improves stand establishment through regulation of apical hook formation and hypocotyl elongation in cotton. Field Crops Res 222, 50–58 (2018).

59. Wang, J. W., Czech, B. & Weigel, D. miR156-Regulated SPL Transcription Factors Define an Endogenous Flowering Pathway in Arabidopsis thaliana. Cell 138, 738–749 (2009).

60. Okushima, Y. et al. Functional Genomic Analysis of the *AUXIN RESPONSE FACTOR* Gene Family Members in *Arabidopsis thaliana*: Unique and Overlapping Functions of *ARF7* and *ARF19* . Plant Cell 17, 444–463 (2005).

61. Ulmasov, T., Murfett, J., Hagen, G. & Guilfoyle, T. J. Aux/IAA proteins repress expression of reporter genes containing natural and highly active synthetic auxin response elements. Plant Cell 9, 1963–1971 (1997).

62. Weigel, D. & Glazebrook, J. Arabidopsis: A Laboratory Manual. (Cold Spring Harbor Laboratory Press, 2022).

63. Patron, N. J. et al. Standards for plant synthetic biology: a common syntax for exchange of DNA parts. New Phytologist 208, 13–19 (2015).

64. Engler, C., Gruetzner, R., Kandzia, R. & Marillonnet, S. Golden Gate Shuffling: A One-Pot DNA Shuffling Method Based on Type IIs Restriction Enzymes. PLoS One 4, 5553 (2009).

65. Clough, S. J. & Bent, A. F. Floral dip: a simplified method for Agrobacterium-mediated transformation of **Arabidopsis thaliana**. The Plant Journal 16, 735– 743 (1998).

