## Supplemental information for "The microRNA156/*SPL9* module mediates auxin response to facilitate apical hook maintenance in *Arabidopsis*"

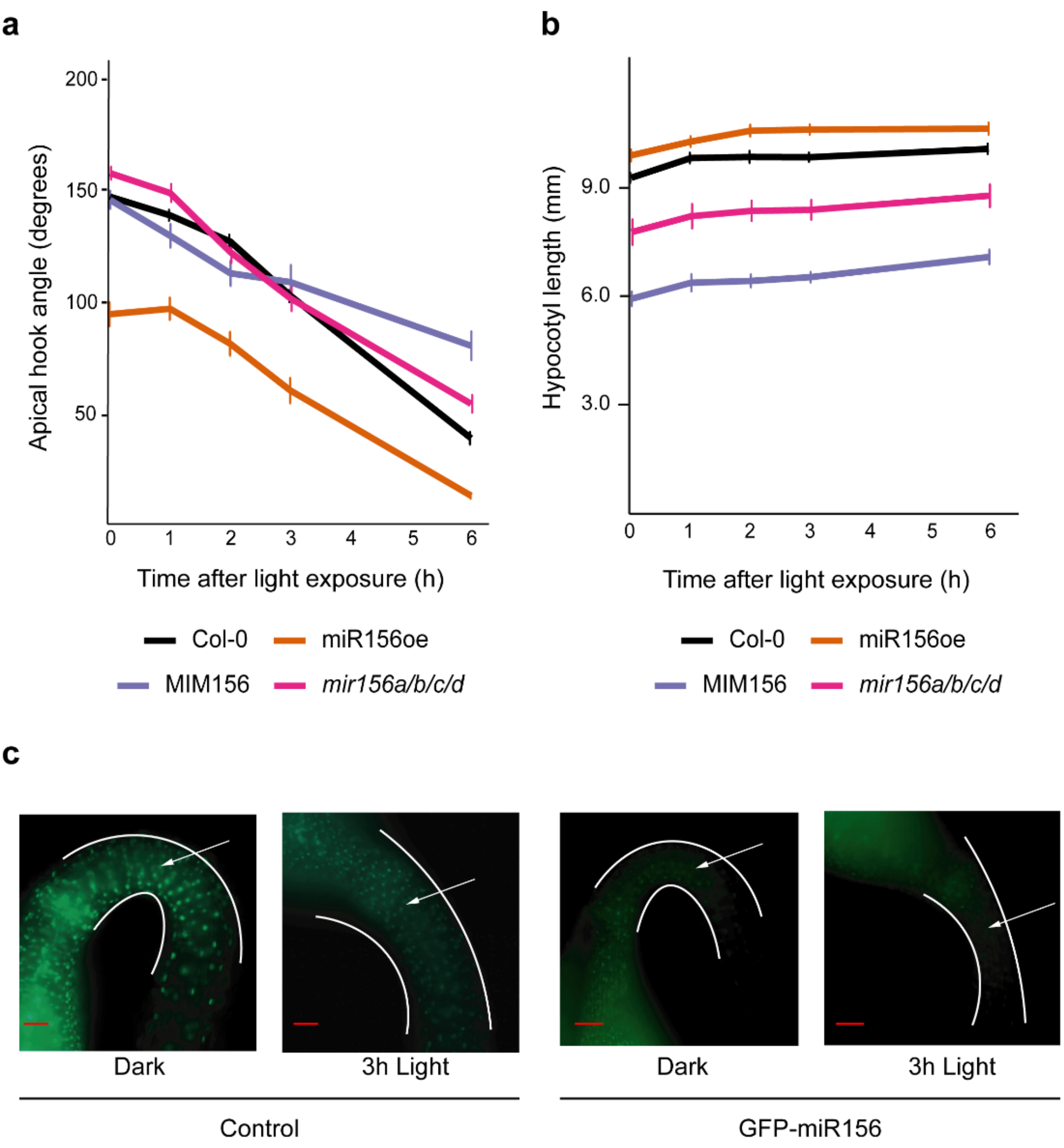


**Supplementary Fig 1. | miR156 regulates apical hook opening. a** Angle of the apical hook of 3-day-old dark-grown seedlings exposed to white light for 1, 2, 3, and 6 hours. Wild type (Col-0), miRNA156 overexpression (miR156oe), Mimicry of miR156 (MIM156), and quadruple loss of miR156 function mutant (*mir156a/b/c/d*). **b** Hypocotyl length of 3-day-old dark-grown seedlings exposed to white light for 1, 2, 3, and 6 hours. Col-0, miR156oe, MIM156, and *mir156a/b/c/d*. **c** Expression of control (*pUBQ10:GFP-N7-TNOS*) and GFP-miR156 (*pUBQ10:GFP-N7-SPL3 3'UTR*) sensors in 3-day-old dark-grown (Dark) seedlings and those exposed to white light for 3 hours (3h Light). Arrows indicate GFP protein accumulation. scale bar: 100 µm.

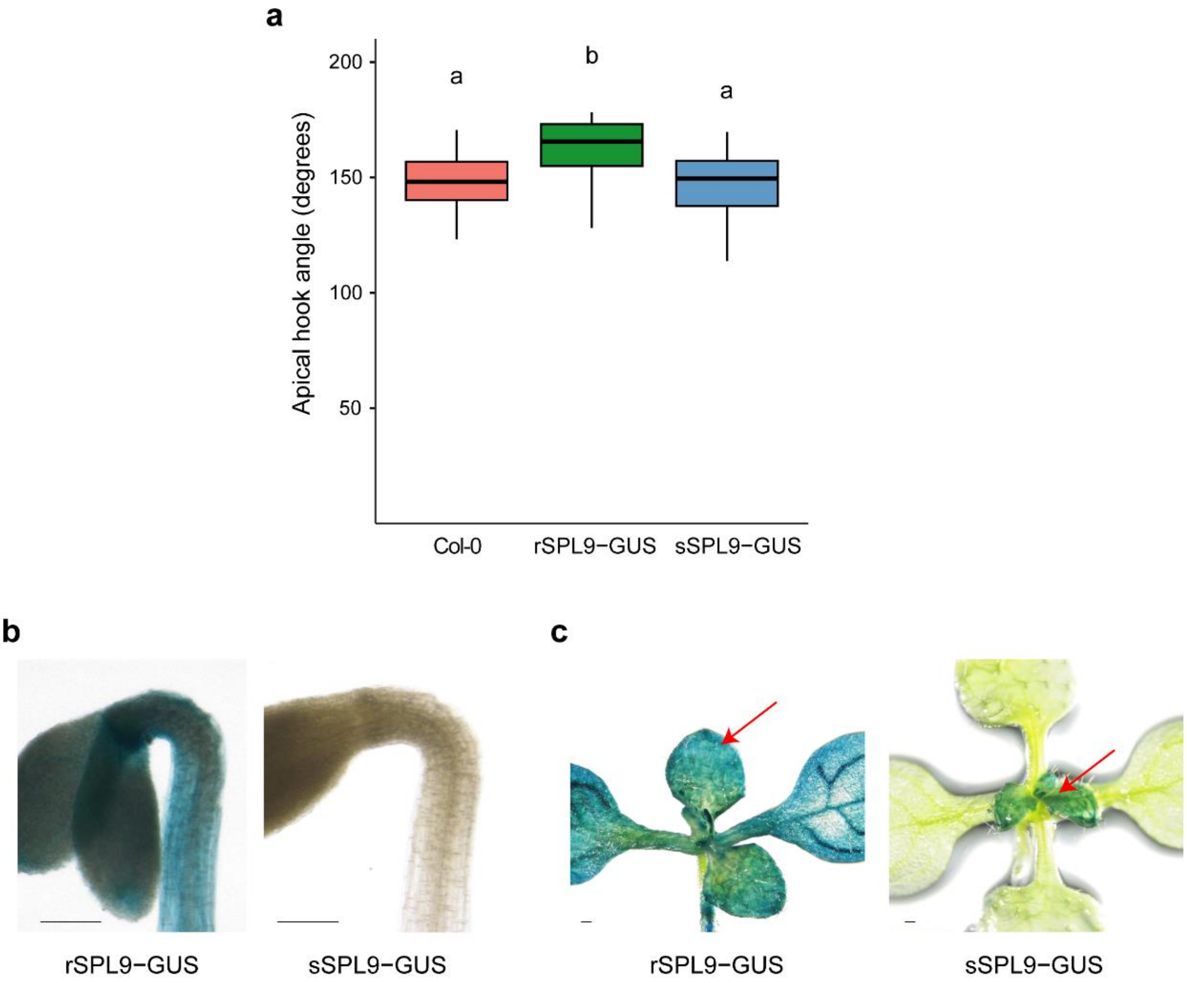


**Supplementary Fig 2. | The miR156-resistant version of *SPL9* increases apical hook angle. a** Apical hook angle of 3-day-old dark-grown seedlings of wild type (Col-0), miR156-resistant version of *SPL9* fused to *GUS* (rSPL9-GUS), and miR156-sensitive version of *SPL9* fused to *GUS* (sSPL9-GUS). Different letters mean statistical significance among the genotypes. ANOVA followed by Tukey test, 1% significance. **b** Representative imagens of 3-day-old dark-grown seedlings showing *GUS* expression. **c** Representative imagens of 10-day-old light-grown seedlings showing *GUS* expression (red arrows). Scale bar: 100 µm.

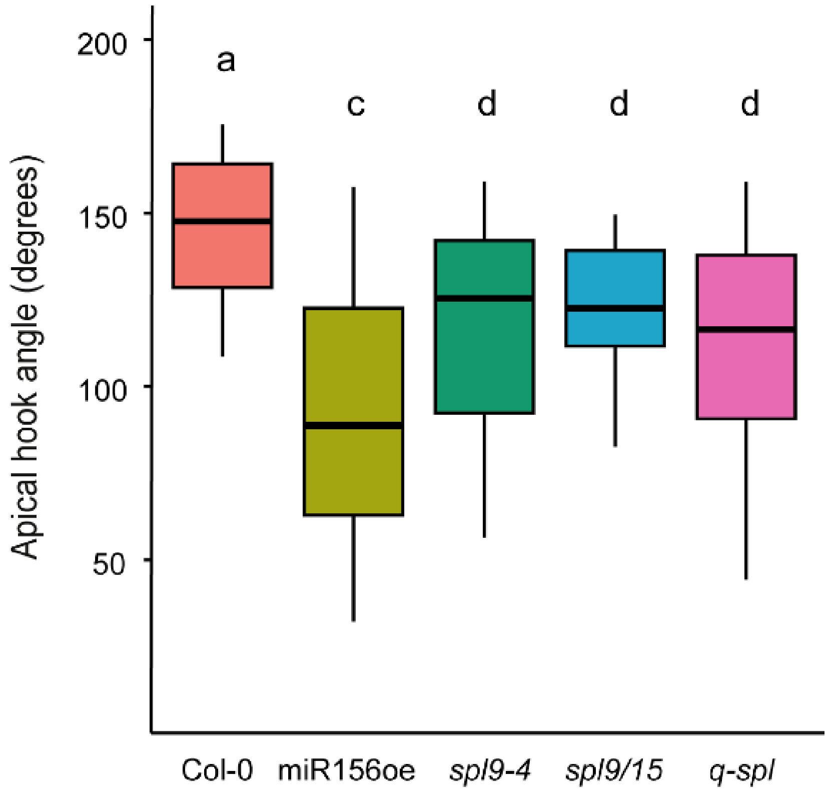


**Supplementary Fig 3. | *SPL9* is required for apical hook development.** Angle of the apical hook of wild type (Col-0); miR156-overexpressing seedling (miR156oe); loss of *SPL9* function (*spl9-4*), loss of *SPL9* and *SPL15* function *(spl9/15)*; and quintuple *spl11 spl13 spl15 spl2 spl9* mutant (*q-spl*). Different letters mean statistical difference among the genotypes. ANOVA followed by Tukey test, 1% significance.

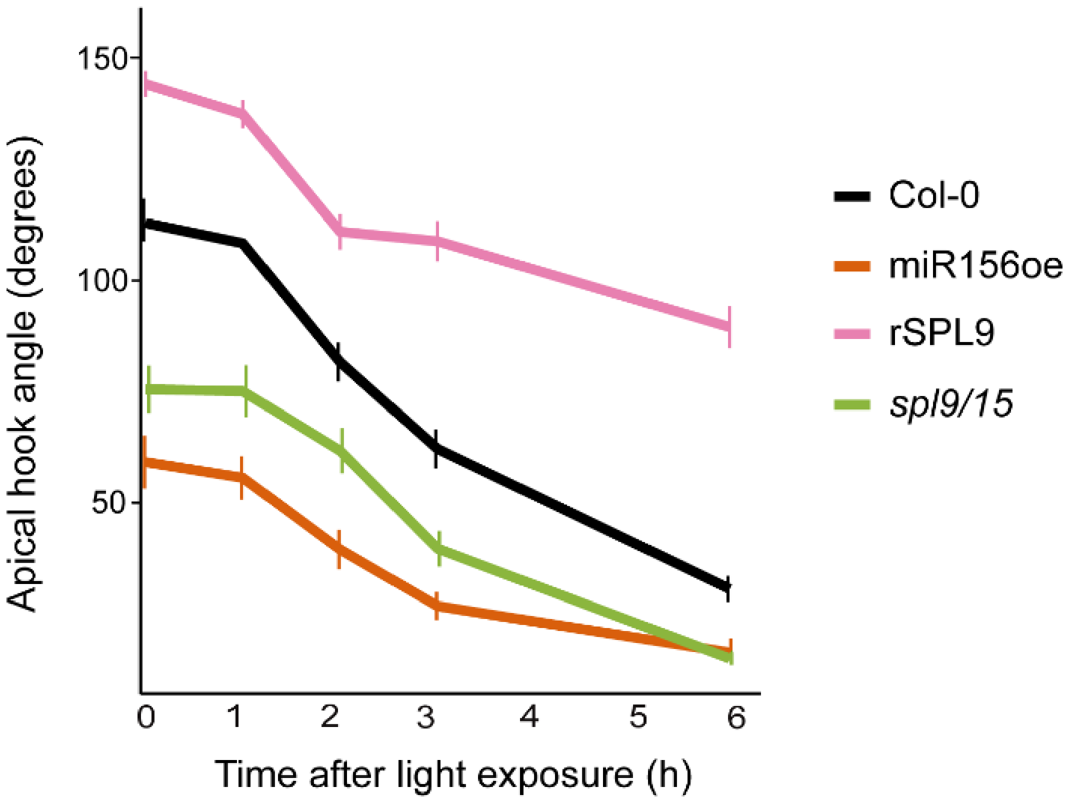


**Supplementary Fig 4. | miR156 and its targets regulate apical hook opening in response to light.** Angle of the apical hook of 3-day-old dark-grown exposed to white light for 1, 2, 3, and 6 hours. Wild type (Col-0), miR156-overexpressing seedling (miR156oe), miR156 resistant version of *SPL9* (rSPL9), and loss of *SPL9* and *SPL15* function *(spl9/15)* mutant.

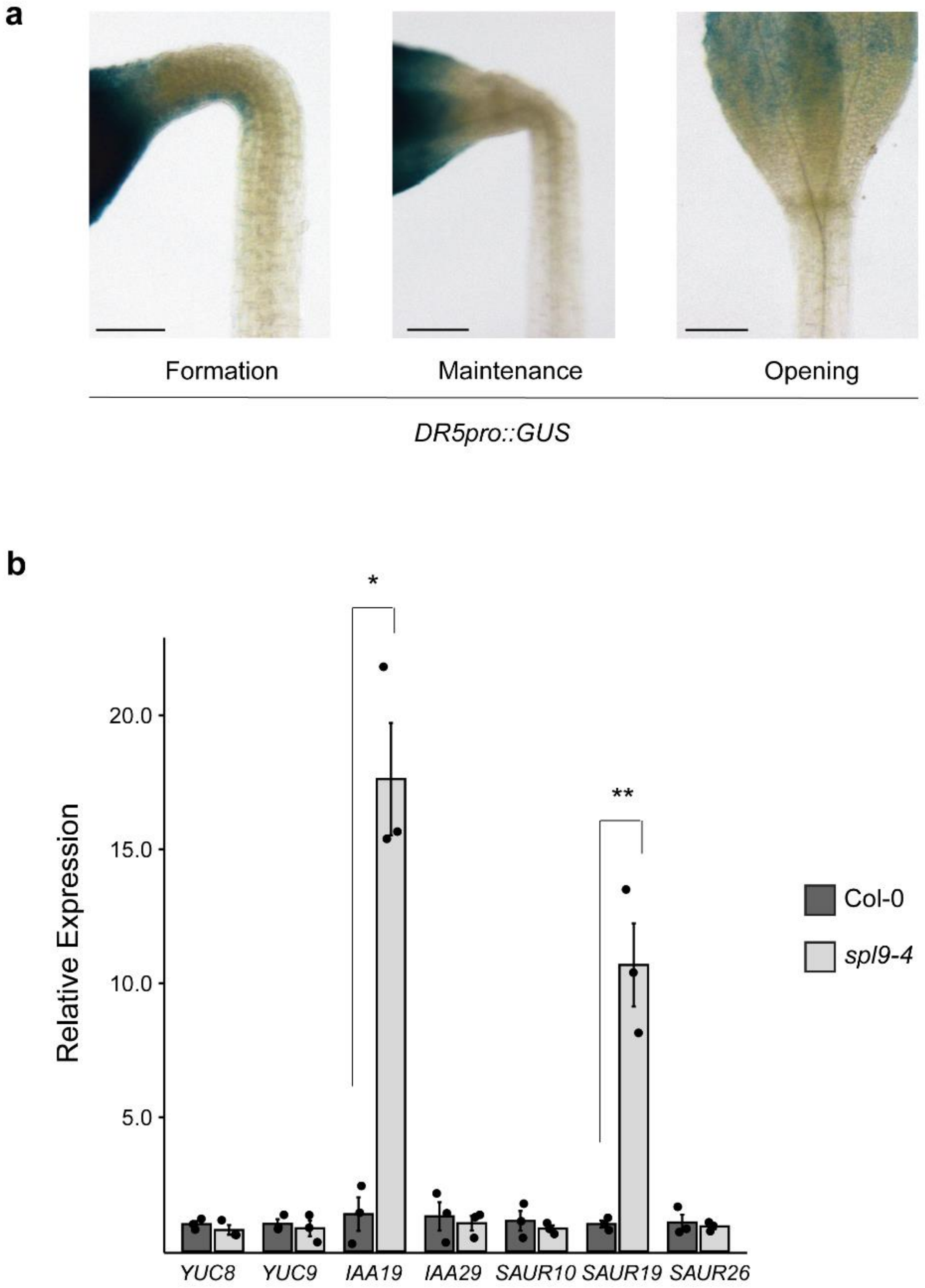


**Supplementary Fig 5. Auxin response in the apical hook is dependent on *SPL9*. a** Expression of *GUS* in *DR5pro::GUS* dark-grown seedlings at formation (2-day-old), maintenance (3-day-old), and opening (4-day-old). **b** Relative expression of *YUCCA8* and *9* (*YUC8* and *YUC9*), *INDOLE-3-ACETIC ACID INDUCIBLE19* and *29* (*IAA19* and *IAA29*), and *small auxin upregulated RNA10, 26,* and *19* (*SAUR10, SAUR19,* and *SAUR26*) in wild type (Col-0) and loss of function of *SPL9* mutant (*spl9-4*). * and **: statistical different by t-test, 5% and 1% significance, respectively.

**Supplementary Table 1.** *Arabidopsis SPL* transcript ratios extracted from the degradome data of de-etiolated seedlings grown from light to dark conditions

| **Gene** | **Light/Dark ratio (Repeat 1)** [a] | **Light/Dark ratio (Repeat 2)** [a] |
|---|---|---|
| *SPL3* | 0.36 | 0.51 |
| ***SPL9*** | **14.83** | **8.84** |
| *SPL2* | 1.48 | 1.06 |
| *SPL13A* | 0.70 | 0.66 |
| *SPL13B* | 0.69 | 0.65 |
| *SPL11* | 1.68 | 1.74 |
| *SPL5* | 1.06 | 0.53 |
| *SPL15* | 2.84 | 2.20 |
| *SPL10* | 1.70 | 1.24 |

[a] miR156 target degradome data obtained from cross comparison of small RNA transcriptome and mRNA degradome of 4-day-old dark-grown seedlings and 4-d-old dark-grown seedlings exposed to white light[20]

**Supplementary Table 2.** Oligonucleotides used in this study.

| Primer ID | Sequence 5’ > 3’ | Target | Application |
|---|---|---|---|
| SPL9pro_F1 F | CGCGTCTCTCTCGGGAGGAGTACTGAATTCATTTATAG | AT2G42200 | Cloning |
| SPL9pro_F1 R | CGCGTCTCTCTCAGAGCAATATTCAGACTCCCC | AT2G42200 | Cloning |
| SPL9pro_F2 F | CGCGTCTCTCTCGGCTCCCTTGTCTCAGATAGA | AT2G42200 | Cloning |
| SPL9pro_F2 R | CGCGTCTCTCTCAATCGTCGAAATTTGTGGAACT | AT2G42200 | Cloning |
| SPL9pro_F3 F | CGCGTCTCTCTCGCGATAGAGACAATACATTTGT | AT2G42200 | Cloning |
| SPL9pro_F3 R | CGCGTCTCTCTCACATTTGACAAGTTAAATGCATTTTCTC | AT2G42200 | Cloning |
| FULpro_F1 F | GGCGTCTCGCTCGGGAGGGTAGTCTAGAGACTTTTCCT | AT5G60910 | Cloning |
| FULpro_F1 R | GGCGTCTCTCTCAGAAGAAGTGACGAGAGTGGAGATAA | AT5G60910 | Cloning |
| FULpro_F2 F | GGCGTCTCGCTCGCTTCCACTCGTATGTAGCTT | AT5G60910 | Cloning |
| FULpro_F2 R | GGCGTCTCTCTCACATTGTATCCTCTCCATGCTAGATT | AT5G60910 | Cloning |
| SAUR19pro_F1 F | GGCGTCTCGCTCGGGAGATGCGAGCCACGTTTGCTTA | AT5G18010 | Cloning |
| SAUR19pro_F1 R | GGCGTCTCTCTCAGAAGAAGGTGTCGTGGCTCAC | AT5G18010 | Cloning |
| SAUR19pro_F2 F | GGCGTCTCGCTCGCTTCAAATATTAAGATCAGGACGT | AT5G18010 | Cloning |
| SAUR19pro_F2 R | GGCGTCTCTCTCACATTATGATAGAAATAATGGGGTTATGG | AT5G18010 | Cloning |
| rSPL9_F1 F | TTGTCGTCTCTCTCGAATGGAGATGGGTTCCAAC | AT2G42200 | Cloning |
| rSPL9_F1 R | CGCGTCTCTCTCAATGCAGGCTGTGGCTTCCTTCGT | AT2G42200 | Cloning |
| rSPL9_F2 F | CGCGTCTCTCTCGGCATCCCTCTCTGTGTTAGCTTCTCGT | AT2G42200 | Cloning |
| rSPL9_F2 R | GCTTAACAAGCTTAAGGCGCAGTTTGAGTCGCCAATTC | AT2G42200 | Cloning |
| rSPL9_F3 F | CGCCTTAAGCTTGTTAAGCAATCCACATCAACCACAT | AT2G42200 | Cloning |
| rSPL9_F3 R | TTGTCGTCTCTCTCAAAGCTCAAAGGGACCAGTTGGT | AT2G42200 | Cloning |
| YUC8 F | GCGTGGACCGCTTGCTGCAACT | AT4G28720 | RT-qPCR |
| YUC8 R | GCGTGGACCGCTTGCTGCAACT | AT4G28720 | RT-qPCR |
| YUC9 F | GCGTGGACCGCTTGCTGCAACT | AT1G04180 | RT-qPCR |
| YUC9 R | GCGTGGACCGCTTGCTGCAACT | AT1G04180 | RT-qPCR |
| IAA19 F | TGGTTCGAGCCAAGGCTATG | AT3G15540 | RT-qPCR |

| | | | |
|---|---|---|---|
| IAA19 R | TCTTTCAAGGCCACACCGAT | AT3G15540 | RT-qPCR |
| IAA29 F | CCCAACGAGGAAGACGAAGA | AT4G32280 | RT-qPCR |
| IAA29 R | TGATCCCACAGTAGCCGTTG | AT4G32280 | RT-qPCR |
| SAUR10 F | GCTTCGTAGGCATTAACCCG | AT2G18010 | RT-qPCR |
| SAUR10 R | CCCCACCCCAACTACAACAA | AT2G18010 | RT-qPCR |
| SAUR19 F | GATTCTAAGCCGCTCCAC | AT5G18010 | RT-qPCR |
| SAUR19 R | CCGAGAAGTCACATTGATG | AT5G18010 | RT-qPCR |
| SAUR26 F | AGTAAAAACAAGCAAGGCACCG | AT3G03850 | RT-qPCR |
| SAUR26 R | CTGCTTCTTCTGGCTCTCGC | AT3G03850 | RT-qPCR |
| FUL F | GAGAAGAAAACGGGTCAGCAAG | AT5G60910 | RT-qPCR |
| FUL R | TGGAGGAGGTTACGCAGTATTGA | AT5G60910 | RT-qPCR |
| ACTIN2 F | GACCTTGCTGGACGTGACCTTAC | AT3G18780 | RT-qPCR |
| ACTIN2 R | GTAGTCAACAGCAACAAAGGAGAGC | AT3G18780 | RT-qPCR |
| spl9-4 LP F | CATGTTTTGCCATTGCCGGA | AT2G42200 | Genotyping |
| spl9-4 RP R | ATGAGGGCAACGGCTTTCTT | AT2G42200 | Genotyping |
| ful-7 LP F | AATTGTCCTTCTTGCTGACCC | AT5G60910 | Genotyping |
| ful-7 RP R | CGATCGAGAAGTTGAGTTTGG | AT5G60910 | Genotyping |
| LBb1 R | GCGTGGACCGCTTGCTGCAACT | Salk T-DNA | Genotyping |
| SPL9pro_gen F | GGTAGATTGGCAATGGACAAACA | AT2G42200 | Genotyping |
| GUS R | TGATACCAGACGTTGCCCG | β-glucuronidase | Genotyping |
